# *Cntnap2* loss drives striatal neuron hyperexcitability and behavioral inflexibility

**DOI:** 10.1101/2024.05.09.593387

**Authors:** Katherine R. Cording, Emilie M. Tu, Hongli Wang, Alexander H. C. W. Agopyan-Miu, Helen S. Bateup

## Abstract

Autism spectrum disorder (ASD) is a neurodevelopmental disorder characterized by two major diagnostic criteria – persistent deficits in social communication and interaction, and the presence of restricted, repetitive patterns of behavior (RRBs). Evidence from both human and animal model studies of ASD suggest that alteration of striatal circuits, which mediate motor learning, action selection, and habit formation, may contribute to the manifestation of RRBs. *CNTNAP2* is a syndromic ASD risk gene, and loss of function of *Cntnap2* in mice is associated with RRBs. How loss of *Cntnap2* impacts striatal neuron function is largely unknown. In this study, we utilized *Cntnap2^−/−^* mice to test whether altered striatal neuron activity contributes to aberrant motor behaviors relevant to ASD. We find that *Cntnap2^−/−^* mice exhibit increased cortical drive of direct pathway striatal projection neurons (dSPNs). This enhanced drive is likely due to increased intrinsic excitability of dSPNs, which make them more responsive to cortical inputs. We find that *Cntnap2^−/−^* mice exhibit spontaneous repetitive behaviors, increased motor routine learning, perseveration, and cognitive inflexibility. Increased corticostriatal drive of the direct pathway may therefore contribute to the acquisition of repetitive, inflexible behaviors in *Cntnap2* mice.

## Introduction

Autism spectrum disorder (ASD) is characterized by alterations in social communication and interaction, as well as the presence of restricted, repetitive, inflexible behaviors (APA, 2022). Given that ASD has high heritability (Sandin et al., 2017), much work has been done in the last 30 years to identify genes that confer risk of developing ASD (De Rubeis et al., 2014; Iossifov et al., 2014; Sanders et al., 2015). Through this, hundreds of high-confidence risk genes have been identified, varying greatly in the proteins for which they code (Satterstrom et al., 2020). These include transcriptional and translational regulators, ion channels, receptors, cell adhesion molecules, and others (De Rubeis et al., 2014; Delorme et al., 2013; Ebert & Greenberg, 2013). Recent work has focused on identifying brain regions and circuits that may be commonly affected by ASD-related mutations. The basal ganglia, in particular the striatum, represents one such commonly altered brain region and prior studies have demonstrated changes in striatal function and striatum-associated behaviors in mice with mutations in ASD risk genes (Benthall et al., 2021; Fuccillo, 2016; Peca et al., 2011; Peixoto et al., 2016; Platt et al., 2017; Rothwell et al., 2014; Wang et al., 2016). However, whether basal ganglia circuits are convergently changed in ASD mouse models is an open question. Here we investigated whether loss of function of the syndromic ASD risk gene *Cntnap2* alters striatal physiology and basal ganglia-dependent behaviors in mice.

*Cntnap2* codes for a neurexin-like cell adhesion molecule called Contactin-associated protein-like 2 (Caspr2) (Poliak et al., 1999; Poliak et al., 2003). In mice, Caspr2 is expressed in several cortical and subcortical regions, including the striatum, from embryonic day 14 (E14) onward into adulthood (Penagarikano et al., 2011). Caspr2 is primarily localized at the juxtaparanodes of myelinated axons and is involved in the clustering of potassium channels (Poliak et al., 2003; Scott et al., 2019). In mice, *in vitro* studies suggests that Caspr2 may also play a role in AMPAR trafficking and cell morphology (Anderson et al., 2012; Gdalyahu et al., 2015; Varea et al., 2015), and *ex vivo* experiments indicate that it can control cell excitability and circuit synchronicity (Martin-de-Saavedra et al., 2022). Caspr2 is important during neurodevelopment and has been implicated in neuronal migration (Penagarikano et al., 2011), the maturation and function of parvalbumin-positive GABAergic interneurons (Penagarikano et al., 2011; Scott et al., 2019; Vogt et al., 2018), and the timing of myelination (Scott et al., 2019). *CNTNAP2* mutations in people lead to a neurodevelopmental syndrome that can include language disorders, epilepsy, obsessive compulsive disorder, and ASD (Penagarikano & Geschwind, 2012; Rodenas-Cuadrado et al., 2014). A mouse model of this syndrome, *Cntnap2^−/−^,* has been shown to exhibit good face validity for ASD-relevant social and motor behavior alterations (Brunner et al., 2015; Dawes et al., 2018; Penagarikano et al., 2011; Scott et al., 2019). However, the impact of *Cntnap2* loss on striatal physiology and corticostriatal-dependent behaviors has not been comprehensively assessed.

The striatum is primarily composed of GABAergic striatal projection neurons (SPNs), which make up two functionally distinct output pathways: the D1-receptor expressing cells of the direct pathway (dSPNs), which project to substantia nigra pars reticulata (SNr), and the D2-receptor expressing cells of the indirect pathway (iSPNs), which project to external globus pallidus (GPe) (Calabresi et al., 2014; Gerfen & Surmeier, 2011; Kravitz et al., 2010; Tai et al., 2012). The two types of SPNs are intermixed throughout the striatum and receive excitatory glutamatergic inputs from cortex and thalamus, as well as dopaminergic input from the midbrain (Ding et al., 2008; Doig et al., 2010; Gerfen & Surmeier, 2011). Coordinated activity between the populations of SPNs in response to these inputs mediates action selection, motor learning and habit formation (Hawes et al., 2015; Santos et al., 2015; Yin et al., 2005, 2006). Although SPNs comprise upwards of 95% of the cells in striatum, there are distinct types of GABAergic interneurons that contribute significantly to the inhibitory circuitry of the striatum. Parvalbumin (PV) interneurons, which make up ∼2% of the cells in striatum, provide the largest feedforward inhibition onto SPNs (Burke et al., 2017). Changes in the number and/or function of PV interneurons have been identified in several ASD mouse models, including *Cntnap2^−/−^* mice, implicating PV circuitry as a potential common alteration across ASD mouse models (Filice et al., 2020; Juarez & Martinez Cerdeno, 2022).

To determine how loss of *Cntnap2* affects striatal function, we assessed the physiology of SPNs and PV-interneurons in the dorsolateral striatum and utilized a range of assays to assess striatum-associated behaviors in *Cntnap2^−/−^* mice. We find that SPNs of the direct pathway exhibit increased corticostriatal drive, despite unchanged excitatory cortical input. Although decreased inhibitory function has been identified in other brain regions in *Cntnap2^−/−^* mice, we find no deficit in broad or PV-specific inhibitory input onto either SPN subtype in the case of *Cntnap2* loss. Instead, we identify a significant increase in the intrinsic excitability of dSPNs in *Cntnap2^−/−^* mice, driven by a reduction in Kv1.2 channel function. Behaviorally, we find that *Cntnap2^−/−^* mice exhibit RRB-like behaviors including increased grooming, marble burying and nose poking in the holeboard assay. *Cntnap2^−/−^* mice also exhibit increased motor routine learning in the accelerating rotarod, and cognitive inflexibility in an odor-based reversal learning task. Taken together, these findings suggest that enhanced direct pathway drive may play a role in the spontaneous and learned repetitive behaviors exhibited by these mice.

## Results

### *Cntnap2^−/−^* dSPNs exhibit increased cortical drive

Emerging evidence indicates that corticostriatal synapses are a common site of alteration in mouse models of ASD (Li & Pozzo-Miller, 2020). To test whether mice with loss of *Cntnap2* exhibit changes in corticostriatal connectivity, we crossed *Cntnap2^−/−^* mice to *Thy1-* ChR2-YFP mice, which express channelrhodopsin in a subset of layer V pyramidal neurons (Fig. 1A) (Arenkiel et al., 2007; Poliak et al., 1999). These mice were crossed to a D1-tdTomato reporter line to visually identify dSPNs (Ade et al., 2011). We recorded from SPNs in the dorsolateral striatum (DLS), as this sensorimotor striatal subregion is implicated in the acquisition of habitual and procedural behaviors (Packard & Knowlton, 2002). Changes in physiological function in this area may be connected to the acquisition of repetitive motor behaviors in ASD (APA, 2022; Fuccillo, 2016). To simulate a train of cortical inputs, we applied ten pulses of blue light over the recording site in DLS and measured the number of action potentials (APs) fired by SPNs in the absence of synaptic blockers (Fig. 1A). We altered the light intensity to vary the probability of eliciting subthreshold depolarizations or AP firing. dSPNs were identified using tdTomato fluorescence, and tdTomato negative neurons were designated putative iSPNs.

**Figure 1.**
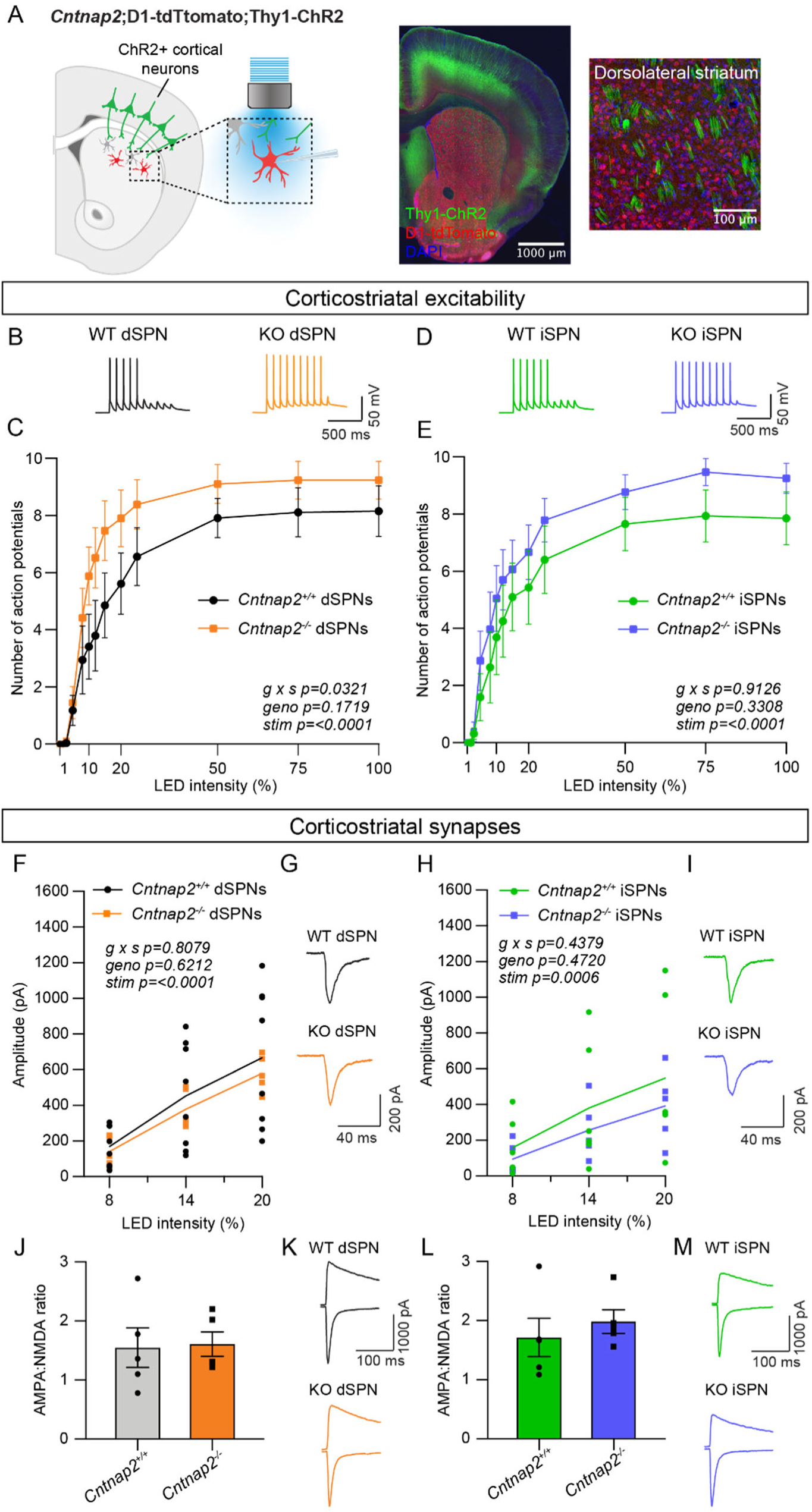
*Cntnap2^−/−^* dSPNs exhibit increased cortical drive. (A) Left: schematic of the corticostriatal connectivity experiments. For corticostriatal excitability, cortical terminals expressing ChR2 were stimulated with 10 pulses of blue light at 10 Hz and responses were recorded from dSPNs (red) and iSPNs (grey) in dorsolateral striatum. For corticostriatal synaptic strength, cortical terminals expressing ChR2 were stimulated with blue light at increasing intensity and synaptic currents were recorded from dSPNs (red) and iSPNs (grey) in dorsolateral striatum. Center: 10x confocal image of the striatum from a *Cntnap2^+/+^;*D1-tdTomato;Thy1-ChR2 mouse. Right: 20x confocal image of dorsolateral striatum from a *Cntnap2^+/+^;*D1-tdTomato;Thy1-ChR2 mouse. YFP (green) labels cell bodies and axons of a subset of layer V pyramidal neurons, tdTomato (red) labels dSPNs, and DAPI stained nuclei are in blue. (B) Example single traces of action potentials (APs) in dSPNs evoked by cortical terminal stimulation at 20% light intensity for the indicated genotypes. (C) Quantification (mean ± SEM) of the number of APs evoked in dSPNs at different light intensities. *Cntnap2^+/+^* n=9 mice, 24 cells, *Cntnap2^−/−^* n=10 mice, 22 cells. Repeated measures two-way ANOVA p values are shown; g x s F (12, 204) = 1.935, geno F (1, 17) = 2.034, stim F (1.931, 32.83) = 86.12. (D) Example single traces of action potentials (APs) in iSPNs evoked by cortical terminal stimulation at 20% light intensity for the indicated genotypes. (E) Quantification (mean ± SEM) of the number of APs evoked in iSPNs at different light intensities. *Cntnap2^+/+^* n=9 mice, 23 cells, *Cntnap2^−/−^* n=10 mice, 21 cells. Repeated measures two-way ANOVA p values are shown; g x s F (12, 216) = 0.5012, geno F (1, 18) = 0.9989, stim F (2.331, 41.96) = 60.62. (F) Average EPSC traces from example dSPNs of each genotype induced by optogenetic cortical terminal stimulation at 14% light intensity. (G) Quantification of EPSC amplitude evoked in dSPNs at different light intensities (line represents the mean, dots/squares represent average EPSC amplitude for each mouse). *Cntnap2^+/+^* n=8 mice, 17 cells, *Cntnap2^−/−^* n=5 mice, 13 cells. Repeated measures two-way ANOVA p values are shown; g x s F (2, 22) = 0.2154, geno F (1, 11) = 0.2585, stim F (1.053, 11.58) = 49.68. (H) Average EPSC traces from example iSPNs of each genotype induced by optogenetic cortical terminal stimulation at 14% light intensity. (I) Quantification of EPSC amplitude evoked in iSPNs at different light intensities (line represents mean, dots/squares represent average for mouse). *Cntnap2^+/+^* n=6 mice, 13 cells, *Cntnap2^−/−^* n=5 mice, 11 cells. Repeated measures two-way ANOVA p values are shown; g x s F (2, 18) = 0.4428, geno F (1, 9) = 0.5635, stim F (1.095, 9.851) = 23.82. (J) Quantification (mean ± SEM) of AMPA:NMDA ratio in dSPNs evoked by 20% light intensity (dots/squares represent average AMPA:NMDA ratio for each mouse). *Cntnap2^+/+^* n=5 mice, 22 cells, *Cntnap2^−/−^* n=5 mice, 22 cells, p=0.8413, Mann-Whitney test. (K) Example traces show pairs of EPSCs evoked by optogenetic corticostriatal stimulation (20% light intensity) recorded at +40 mV (top traces) and −70 mV (bottom traces) from *Cntnap2^+/+^ and Cntnap2^−/−^* dSPNs. (L) Quantification (mean ± SEM) of AMPA:NMDA ratio in iSPNs evoked by 20% light intensity (dots/squares represent average AMPA:NMDA ratio for each mouse). *Cntnap2^+/+^* n=5 mice, 21 cells, *Cntnap2^−/−^* n=5 mice, 21 cells, p=0.3095, Mann-Whitney test. (M) Example traces show pairs of EPSCs evoked by optogenetic corticostriatal stimulation (20% light intensity) recorded at +40 mV (top traces) and −70 mV (bottom traces) from *Cntnap2^+/+^ and Cntnap2^−/−^* iSPNs.

We quantified the number of evoked APs at different light intensities and found that dSPNs in young adult *Cntnap2^−/−^* mice exhibited increased spike probability compared to wild-type (WT) dSPNs (Fig. 1B-C). The interaction effect of genotype and stimulation intensity in these cells suggests increased corticostriatal drive, consistent with findings in another mouse model with loss of function of the ASD-risk gene *Tsc1* (Benthall et al., 2021). *Cntnap2^−/−^* iSPNs had subtly increased cortically-evoked AP firing compared to WT iSPNs, although this was not statistically significant (1D-E). To test whether the enhanced spiking probability of *Cntnap2^−/−^* dSPNs was due to excitatory synaptic changes, we applied blue light of varying intensity over the recording site in DLS while holding cells at –70 mV to evoke AMPAR-driven excitatory postsynaptic currents (EPSCs). We found that the average optically-evoked EPSC amplitude was not significantly different across a range of light intensities in *Cntnap2^−/−^* dSPNs or iSPNs compared to WT (Fig. 1F-I). In an additional group of mice, we measured the ratio of AMPAR currents recorded at −70 mV to NMDAR currents recorded at +40 mV (at 20% blue light intensity) and found no significant differences in AMPA:NMDA ratio in *Cntnap2^−/−^* dSPNs or iSPNs (Fig. 1J-M).

To further assess synaptic inputs, we measured the number of dendritic spines in *Cntnap2^−/−^* and WT SPNs, which are typically the sites of cortical synaptic innervation (Bouyer et al., 1984; Xu et al., 1989). To visualize spines, we injected neonate *Cntnap2;Drd1a*-tdTomato mice with AAV5-*Syn1*-GFP virus to sparsely label dSPNs and iSPNs in the DLS (Fig. S1) (Keaveney et al., 2018). We found that *Cntnap2^−/−^* SPNs in adult mice had similar spine density as WT (Fig. S1), suggesting no overall change in synapse number. Together, these results show that dSPNs in *Cntnap2^−/−^* mice exhibit enhanced cortically-driven spiking. However, this is not due to a change in corticostriatal synaptic strength or overall synapse density.

### Inhibition is unchanged in *Cntnap2^−/−^* SPNs

Previous work has indicated a reduction in the number and/or function of fast-spiking parvalbumin-expressing (PV) interneurons across multiple brain regions in *Cntnap2^−/−^* mice (Ahmed et al., 2023; Antoine et al., 2019; Jurgensen & Castillo, 2015; Paterno et al., 2021; Penagarikano et al., 2011; Vogt et al., 2018). While inhibitory deficits have been identified in the cortex and hippocampus (Antoine et al., 2019; Jurgensen & Castillo, 2015), and the number of PV interneurons has been reported to be decreased in striatum (Penagarikano et al., 2011), a comprehensive assessment of inhibitory synaptic function has yet to be completed in the striatum of *Cntnap2^−/−^* mice. We first determined whether there were global deficits in inhibition onto SPNs in *Cntnap2^−/−^* mice using intrastriatal electrical stimulation to evoke inhibitory postsynaptic currents (IPSCs) (Fig. 2A). In *Cntnap2^−/−^* dSPNs and iSPNs, the average amplitude of electrically-evoked IPSCs across a range of stimulation intensities was not different from WT (Fig. 2B-E).

**Figure 2.**
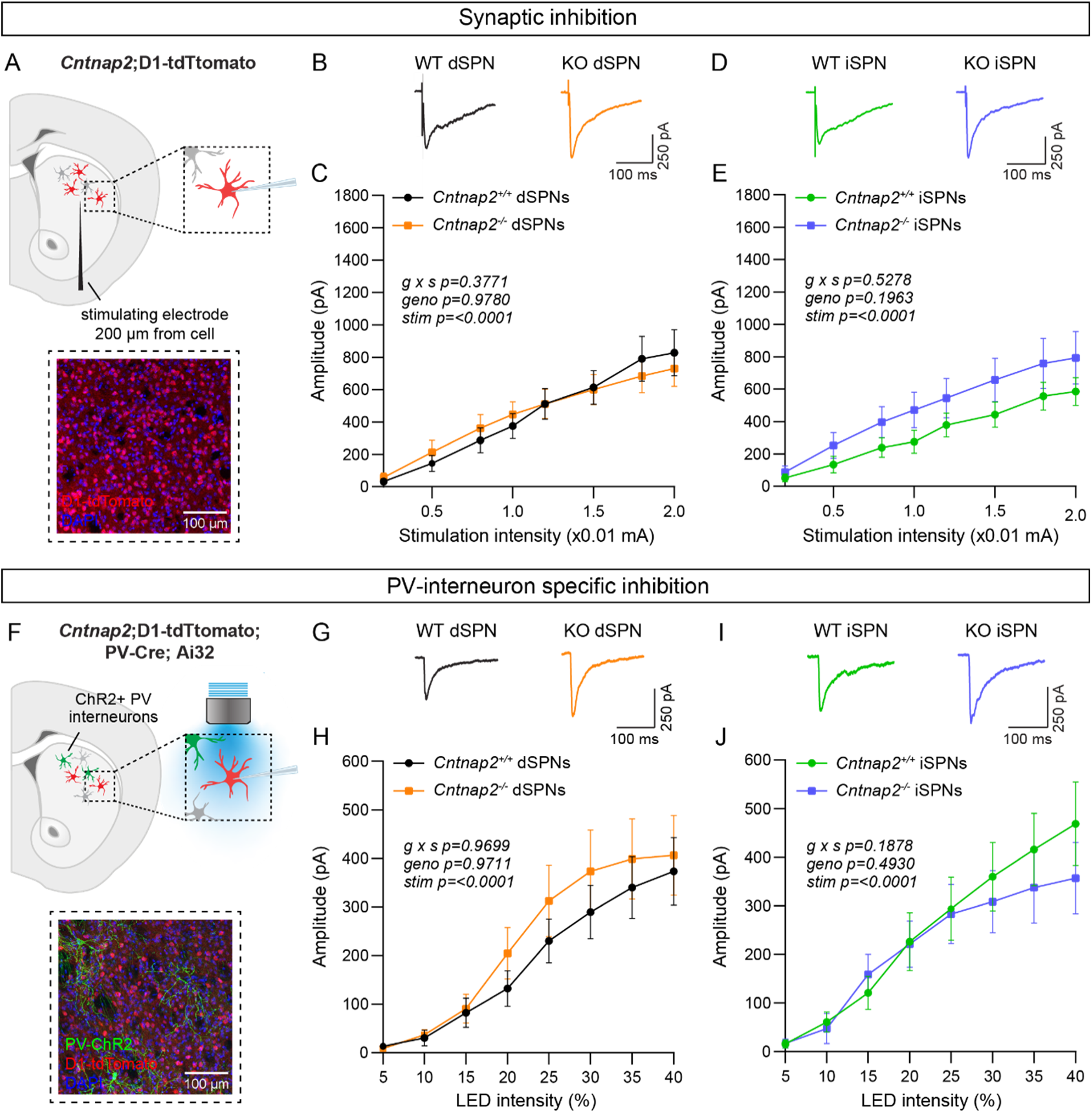
Inhibition is not altered in *Cntnap2^−/−^* SPNs. (A) Top: schematic of the experiment. A bipolar stimulating electrode was placed approximately 200 μm from the recording site. A range of electrical stimulation intensities was applied to the tissue while IPSCs were recorded from dSPNs (red) and iSPNs (grey) in dorsolateral striatum. Bottom: 20x confocal image of dorsolateral striatum from a *Cntnap2^+/+^;*D1-tdTomato mouse. tdTomato (red) labels dSPNs, DAPI stained nuclei are in blue. (B) Average IPSC traces from example dSPNs of each genotype evoked by electrical stimulation at 1.5 (x0.01 mA) intensity for the indicated genotypes. (C) Quantification (mean ± SEM) of IPSC amplitude in dSPNs at different stimulation intensities. *Cntnap2^+/+^* n=17 cells from 9 mice, *Cntnap2^−/−^* n=16 cells from 9 mice. Repeated measures two-way ANOVA p values are shown; g x s F (7, 217) = 1.080, geno F (1, 31) = 0.0007751, stim F (1.815, 56.28) = 54.92. (D) Average IPSC traces from example iSPNs of each genotype evoked by electrical stimulation at 1.5 (x0.01 mA) intensity for the indicated genotypes. (E) Quantification (mean ± SEM) of IPSC amplitude in iSPNs at different stimulation intensities. *Cntnap2^+/+^* n=16 cells from 9 mice, *Cntnap2^−/−^* n=16 cells from 10 mice. Repeated measures two-way ANOVA p values are shown; g x s F (7, 210) = 0.8741, geno F (1, 30) = 1.746, stim F (1.591, 47.73) = 45.66. (F) Top: schematic of the experiment. PV interneuron terminals expressing ChR2 were stimulated with blue light at a range of intensities and optically-evoked IPSCs were recorded from dSPNs (red) and iSPNs (grey) in dorsolateral striatum. Bottom: 20x confocal image of dorsolateral striatum from a *Cntnap2^+/+^;*D1-tdTomato;PV-Cre;Ai32 mouse. YFP (green) labels PV interneurons, tdTomato (red) labels dSPNs, DAPI stained nuclei are in blue. (G) Average IPSC traces from example dSPNs of each genotype evoked by optogenetic PV interneuron stimulation at 30% light intensity. (H) Quantification (mean ± SEM) of IPSC amplitude in dSPNs at different light intensities. *Cntnap2^+/+^* n=29 cells from 15 mice, *Cntnap2^−/−^* n=23 cells from 11 mice. Repeated measures two-way ANOVA p values are shown; g x s F (7, 441) = 0.2566, geno F (1, 63) = 0.001322, stim F (1.433, 90.25) = 32.57. (I) Average IPSC traces from example iSPNs of each genotype evoked by optogenetic PV interneuron stimulation at 30% light intensity. (J) Quantification (mean ± SEM) of IPSC amplitude in iSPNs at different light intensities. *Cntnap2^+/+^* n=24 cells from 14 mice, *Cntnap2^−/−^* n=27 cells from 13 mice. Repeated measures two-way ANOVA p values are shown; g x s F (7, 343) = 1.441, geno F (1, 49) = 0.4771, stim F (1.622, 79.46) = 38.49.

There are many sources of inhibition in the striatum (Burke et al., 2017), which can all be activated with electrical stimulation. To assess whether inhibition from PV interneurons specifically is altered in *Cntnap2^−/−^* mice, we crossed *Cntnap2^−/−^*;D1-tdTomato mice to *Pvalb*-Cre; RCL-ChR2-H134R-EYFP (Ai32) mice to express channelrhodopsin in PV interneurons (Fig. 2F) (Hippenmeyer et al., 2005; Madisen et al., 2010). We applied a blue light pulse of varying intensity over the recording site to evoke PV interneuron-specific IPSCs in SPNs, in the presence of excitatory synaptic blockers (Fig. 2F). We found that the average amplitude of optically-evoked IPSCs did not differ significantly in *Cntnap2^−/−^* dSPNs or iSPNs compared to WT controls (Fig. 2G-J).

To directly measure PV neuron function, we assessed the intrinsic excitability of PV interneurons in *Cntnap2^−/−^* mice. To visualize PV interneurons for recordings, we crossed *Cntnap2^−/−^* mice to *Pvalb*-Cre;RCL-tdT (Ai9) mice (Fig. S2A). Plotting the number of APs fired as a function of current step amplitude indicated that there were no significant differences in the intrinsic excitability of PV interneurons in *Cntnap2^−/−^* mice compared to controls (Fig. S2B,C). There were also no changes in intrinsic cell properties such as rheobase, membrane resistance, capacitance, resting membrane potential, or AP shape in *Cntnap2^−/−^* PV interneurons (Fig. S2D-K).

Given prior reports of altered PV cell number in *Cntnap2^−/−^* mice (Paterno et al., 2021; Penagarikano et al., 2011; Vogt et al., 2018), we counted PV-expressing cells in the striatum, using immunohistochemistry and fluorescent in situ hybridization. We found no significant difference in the number of PV-positive cells in the dorsal striatum of *Cntnap2^−/−^* mice compared to WT (Fig. S3A-F and K-N). We also found no changes in the average single cell or total level of PV protein expression in the dorsal striatum (Fig. 3G-J). Overall, we did not observe significant changes in PV interneuron number, PV expression, or PV interneuron-mediated inhibition in the adult *Cntnap2^−/−^* striatum compared to WT controls.

**Figure 3.**
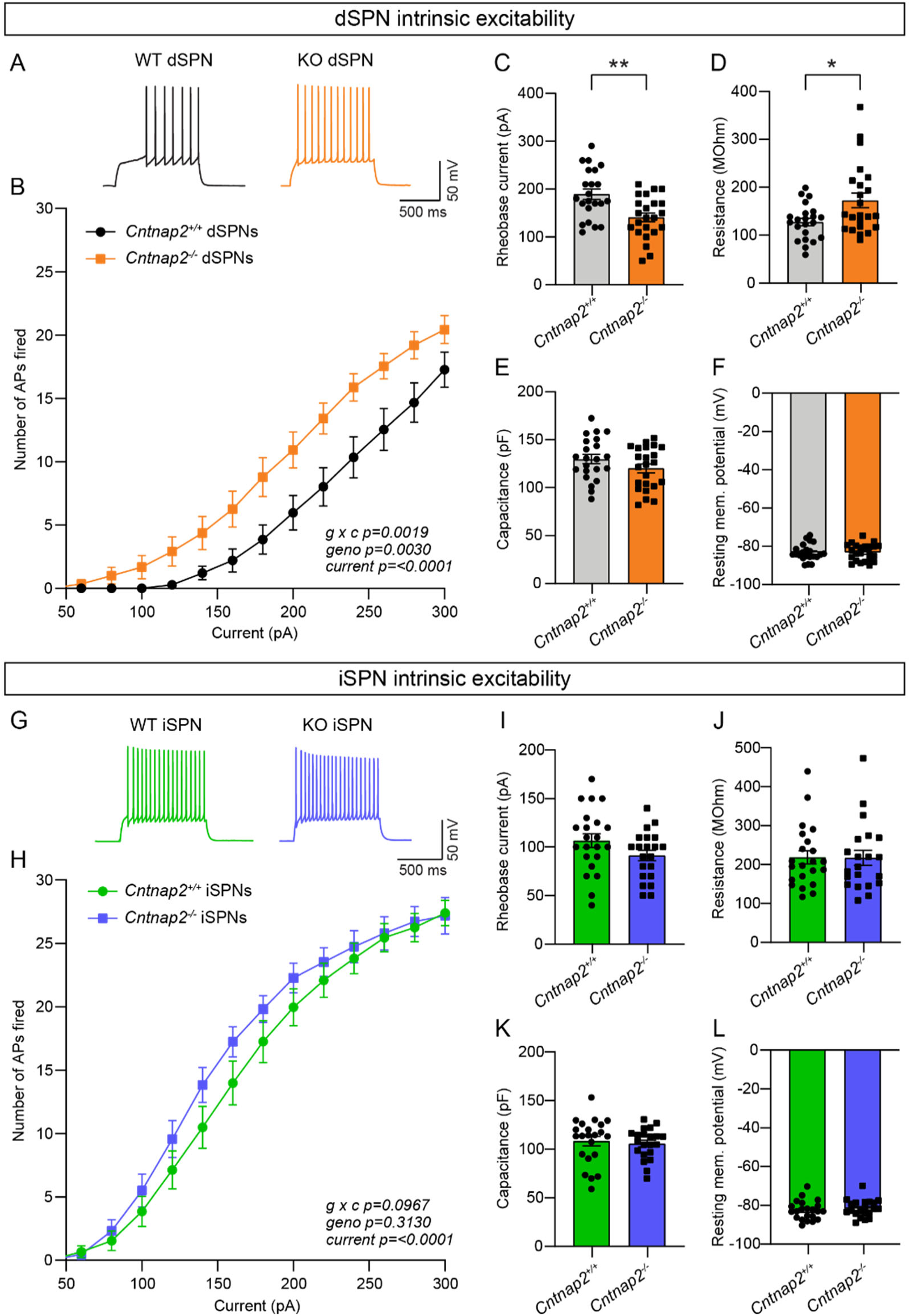
Intrinsic excitability is increased in *Cntnap2^−/−^* dSPNs. (A) Example AP traces in dSPNs evoked by a 200 pA current step for the indicated genotypes. (B) Quantification (mean ± SEM) of the number of APs evoked in dSPNs at different current step amplitudes. *Cntnap2^+/+^* n=22 cells from 8 mice, *Cntnap2^−/−^* n=23 cells from 8 mice. Repeated measures two-way ANOVA p values are shown; g x c F (12, 528) = 2.649, geno F (1, 44) = 107.5, current F (1.974, 86.86) = 147.5. (C) Quantification (mean ± SEM) of the rheobase current in dSPNs. Dots/squares represent the rheobase current for each neuron. n is the same as in panel B. **p=0.0016, two-tailed unpaired t test. (D-F) Quantification (mean ± SEM) of dSPN membrane resistance (D), *p=0.0328, Mann-Whitney test; membrane capacitance (E), p=0.2182, Mann-Whitney test; and resting membrane potential (F), p=0.9914, two-tailed unpaired t test. Dots/squares represent the average value for each neuron. n is the same as in panel B. (G) Example AP traces in iSPNs evoked by a 200 pA current step for the indicated genotypes. (H) Quantification (mean ± SEM) of the number of APs evoked in iSPNs at different current step amplitudes. *Cntnap2^+/+^* n=22 cells from 8 mice, *Cntnap2^−/−^* n=21 cells from 8 mice. Repeated measures two-way ANOVA p values are shown; g x c F (12, 516) = 1.569, geno F (1, 43) = 1.042, current F (2.041, 87.78) = 284.7. (I) Quantification (mean ± SEM) of the rheobase current in iSPNs. Dots/squares represent the rheobase current for each neuron. n is the same as in panel H. p=0.0923, two-tailed unpaired t test. (J-L) Quantification (mean ± SEM) of iSPN membrane resistance (J), p=0.8193, Mann-Whitney test; membrane capacitance (K), p=0.6886, two-tailed unpaired t test; and resting membrane potential (L), p=0.4859, two-tailed unpaired t test. Dots/squares represent the average value for each neuron. n is the same as in panel H.

### dSPN intrinsic excitability is increased in *Cntnap2^−/−^* mice

Given that the increased cortical drive onto *Cntnap2^−/−^* dSPNs could not be explained by changes in excitatory or inhibitory synaptic function, we tested whether it could be due to a change in intrinsic excitability. To measure this, we recorded from dSPNs and iSPNs in *Cntnap2*;D1-tdTomato mice and injected current steps of increasing amplitude. We found that *Cntnap2^−/−^* dSPNs had significantly increased intrinsic excitability compared to WT dSPNs (Fig. 3A-B). *Cntnap2^−/−^* dSPNs also had reduced rheobase current (Fig. 3C), the minimum current required to evoke an AP, as well as increased membrane resistance (Fig. 3D). While there was a trend towards increased excitability in *Cntnap2^−/−^* iSPNs, this effect was not statistically significant (Fig. 3G-H), and these cells did not exhibit changes in rheobase current (Fig. 3I) or membrane resistance (Fig. 3J). Membrane capacitance (Fig. 3E,K), resting membrane potential (Fig. 3F,L), and AP shape (Fig. S4) were not significantly changed in *Cntnap2^−/−^* SPNs, although latency to first spike was reduced in both *Cntnap2^−/−^* dSPNs and iSPNs (Fig. S4C, I). Given the lack of synaptic changes observed in *Cntnap2^−/−^* dSPNs, the increase in dSPN intrinsic excitability likely underlies their enhanced corticostriatal drive (see Fig. 1).

### The effects of Kv1.2 blockade are occluded in *Cntnap2^−/−^* dSPNs

Caspr2 is known to be involved in the clustering of voltage-gated potassium channels (Inda et al., 2006; Poliak et al., 1999; Poliak et al., 2003), particularly Kv1.2 channels (Scott et al., 2019). These channels play an important role in regulating the intrinsic excitability of SPNs (Nisenbaum et al., 1994). Blockade of these channels with the drug α-Dendrotoxin (α-DTX), results in increased AP frequency, decreased rheobase current, decreased first AP latency, and decreased AP threshold (Shen et al., 2004), particularly in dSPNs (Lahiri & Bevan, 2020). Given that *Cntnap2^−/−^* dSPNs exhibited increased AP frequency (Fig. 3B), decreased rheobase current (Fig. 3C), decreased first AP latency (Fig. S4C), and a trend towards decreased AP threshold (p=0.0516; Fig. S4B), we hypothesized that loss of function of Kv1.2 channels could be the mechanism. To test this, we recorded from dSPNs and iSPNs in *Cntnap2*;*Drd1a*-tdTomato mice and injected current steps of increasing amplitude in the absence or presence of α-DTX (100nM). We found that α-DTX enhanced AP firing and decreased AP threshold in WT dSPNs (Fig. 4A-C), but not in *Cntnap2^−/−^* dSPNs (Fig. 4D-F). AP width was also significantly increased by α-DTX in WT but not *Cntnap2^−/−^* dSPNs (Fig. S5H,Q), while AP fast afterhyperpolarization and AP latency were significantly decreased in both WT and *Cntnap2^−/−^* dSPNs (Fig. S5E, I, N, R). Together these data show that the effects of α-DTX on *Cntnap2^−/−^* dSPNs are largely occluded, indicating that Kv1.2 channels are basally altered by loss of *Cntnap2*. This likely accounts for the enhanced intrinsic excitability of *Cntnap2^−/−^* dSPNs.

**Figure 4.**
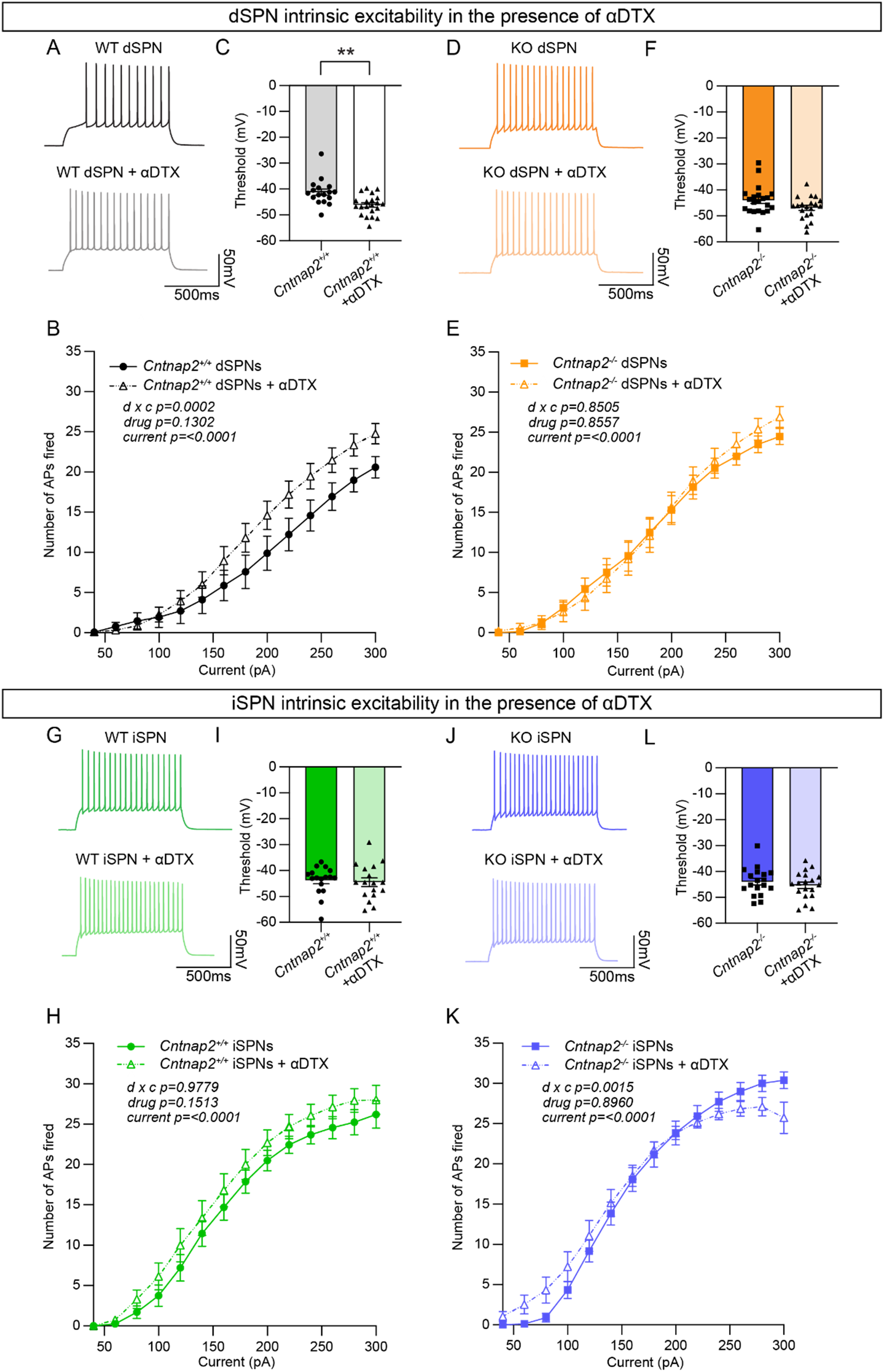
The effects of Kv1.2 blockade are occluded in *Cntnap2^−/−^* dSPNs. (A) Example AP traces from *Cntnap2^+/+^* dSPNs evoked by a 200 pA current step in the absence or presence of α-Dendrotoxin (α-DTX). (B) Quantification (mean ± SEM) of the number of APs evoked in *Cntnap2^+/+^* dSPNs in the absence or presence of α-DTX at different current step amplitudes. *Cntnap2^+/+^* n=17 cells from 8 mice, *Cntnap2^+/+^* + α-DTX n=21 cells from 8 mice. Repeated measures two-way ANOVA p values are shown; drug x c F (13, 468) = 3.097, drug F (1, 36) = 2.399, current F (2.091, 75.28) = 172.0. (C) Quantification (mean ± SEM) of the AP threshold in *Cntnap2^+/+^* dSPNs in the absence or presence of α-DTX. Dots/triangles represent the threshold for each neuron. n is the same as in panel B. **p=0.0010, Mann-Whitney test. (D) Example AP traces from *Cntnap2^−/−^* dSPNs evoked by a 200 pA current step in the absence or presence of α-DTX. (E) Quantification (mean ± SEM) of the number of APs evoked in *Cntnap2^−/−^* dSPNs. *Cntnap2^−/−^* n=21 cells from 9 mice, *Cntnap2^−/−^* + α-DTX n=20 cells from 9 mice. Repeated measures two-way ANOVA p values are shown; drug x c F (13, 507) = 0.6054, drug F (1, 39) = 0.03352, current F (2.156, 84.07) = 247.8. (F) Quantification (mean ± SEM) of the AP threshold in *Cntnap2^−/−^* dSPNs. Squares/triangles represent the threshold for each neuron. n is the same as in panel E. p=0.1348, Mann-Whitney test. (G) Example AP traces in *Cntnap2^+/+^* iSPNs evoked by a 200 pA current step in the absence or presence of α-DTX. (H) Quantification (mean ± SEM) of the number of APs evoked in *Cntnap2^+/+^* iSPNs. *Cntnap2^+/+^* n=17 cells from 9 mice, *Cntnap2^+/+^* + α-DTX n=17 cells from 9 mice. Repeated measures two-way ANOVA p values are shown; drug x c F (13, 416) = 0.3719, drug F (1, 32) = 2.161, current F (1.955, 62.57) = 215.9. (I) Quantification (mean ± SEM) of the AP threshold in *Cntnap2^+/+^* iSPNs. Dots/triangles represent the threshold for each neuron. n is the same as in panel H. p=0.5401, Mann-Whitney test. (J) Example AP traces in *Cntnap2^−/−^* iSPNs evoked by a 200 pA current step in the absence or presence of α-DTX. (K) Quantification (mean ± SEM) of the number of APs evoked in *Cntnap2^−/−^* iSPNs. *Cntnap2^−/−^* n=18 cells from 10 mice, *Cntnap2^−/−^* + α-DTX n=19 cells from 10 mice. Repeated measures two-way ANOVA p values are shown; drug x c F (13, 455) = 2.623, drug F (1, 35) = 0.01734, current F (1.721, 60.23) = 227.4. (L) Quantification (mean ± SEM) of the AP threshold in *Cntnap2^−/−^* iSPNs. Squares/triangles represent the threshold for each neuron. n is the same as in panel K. p=0.4250, two-tailed unpaired t test.

We tested the effects of α-DTX on WT and *Cntnap2^−/−^* iSPNs and found minimal changes (Fig. 4G-K, Fig. S6), with no alterations in AP threshold (Fig. 4I,L) and only a small shift in the input-output relationship of *Cntnap2^−/−^* iSPNs (Fig. 4K). The effects of α-DTX on iSPN excitability have been less well documented and our data suggest a difference in the sensitivity of dSPNs and iSPNs to Kv1 blockade.

### *Cntnap2^−/−^* mice display increased repetitive behaviors

RRBs comprise one of the primary symptom domains of ASD (APA, 2022). Alterations in striatal circuits are thought to be involved in the manifestation of RRBs, given the striatum’s role in action selection and motor control (Estes et al., 2011; Fuccillo, 2016; Hollander et al., 2005; Langen et al., 2014). To determine whether changes in motor behavior accompanied the altered striatal physiology in *Cntnap2^−/−^* mice, we assessed locomotor activity and spontaneous repetitive behaviors using the open field, marble burying, and holeboard assays (Fig. 5A-D). In the open field, we found no significant difference in the total distance traveled, average speed, or number of rears in *Cntnap2^−/−^* mice compared to WT controls (Fig. 5E-G). We did find that *Cntnap2^−/−^* mice made significantly more entries into the center of the open field arena than WT mice, which may reflect a reduction in avoidance behavior in these mice (Fig. 5H). Manually scored grooming behavior revealed that *Cntnap2^−/−^* mice initiated more grooming bouts than WT controls in the open field (Fig. 5I), consistent with prior reports (Penagarikano et al., 2011).

**Figure 5.**
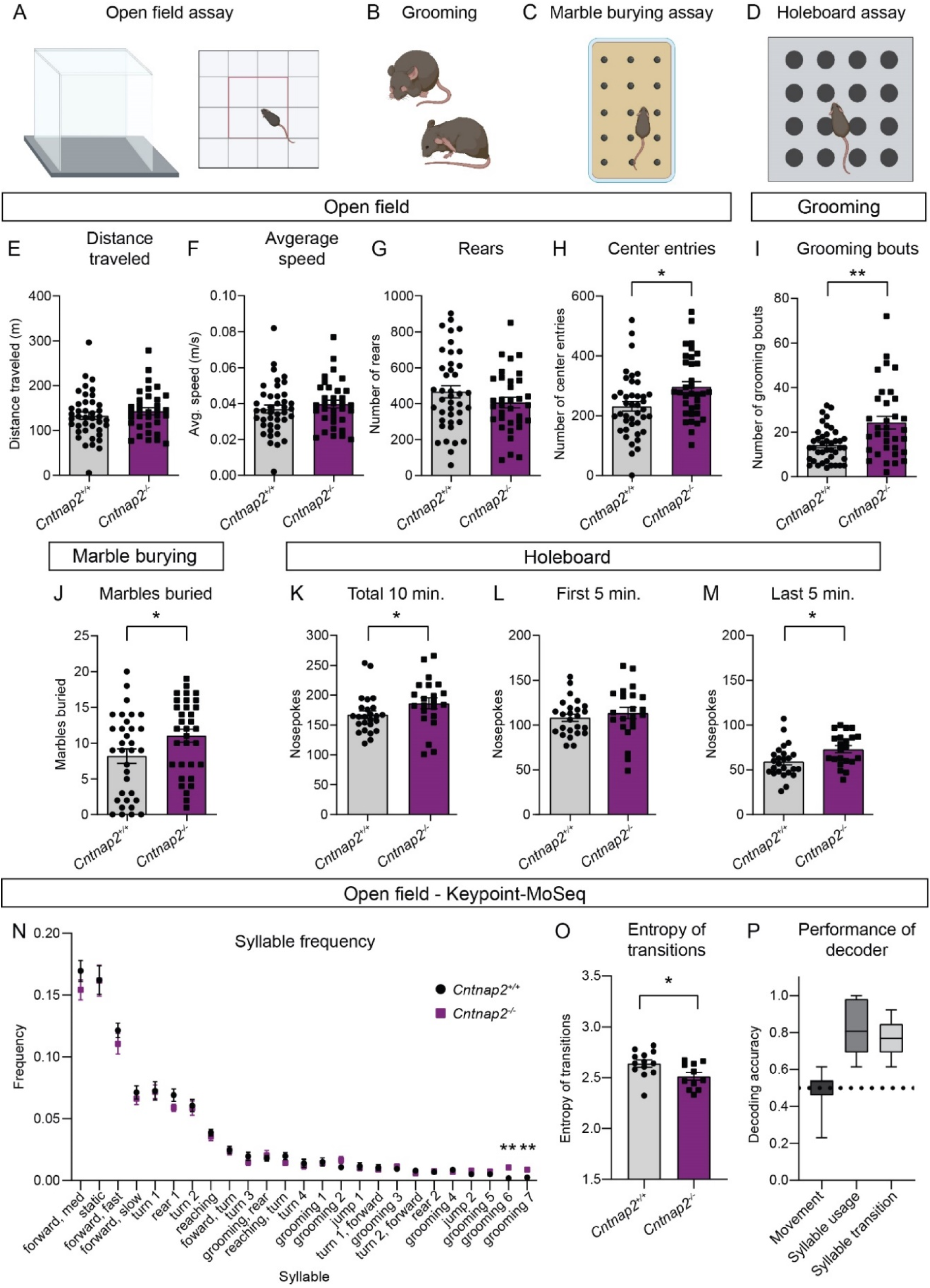
*Cntnap2^−/−^* mice have increased repetitive behaviors. (A-D) Schematics of the behavioral assays used to measure repetitive behaviors in *Cntnap2^+/+^* and *Cntnap2^−/−^* mice, created with BioRender.com. Male and female mice were used for all tests. (E-H) Quantification (mean ± SEM) of open field activity over 60 minutes. (E) total distance traveled, p= 0.3538, Mann-Whitney test; (F) average speed, p=0.3832, Mann-Whitney test; (G) number of rears, p=0.1892, two-tailed unpaired t test; (H) number of center entries, *p=0.0101, two-tailed unpaired t test. *Cntnap2^+/+^* n=41 mice, *Cntnap2^−/−^* n=34 mice. (I) Quantification (mean ± SEM) of the number of manually scored grooming bouts in the first 20 minutes of the open field test, **p=0.0034, Mann-Whitney test, *Cntnap2^+/+^* n=41 mice, *Cntnap2^−/−^* n=34 mice. (J) Quantification (mean ± SEM) of total marbles buried in the marble burying assay. *Cntnap2^+/+^* n=33 mice and *Cntnap2^−/−^* n=33 mice, *p=0.0396, two-tailed unpaired t test. (K-M) Quantification (mean ± SEM) of performance in the holeboard assay. (K) Total number of nose pokes in 10 minutes, *p=0.0212, Mann-Whitney test; (L) nose pokes in the first five minutes, p=0.4811, two-tailed unpaired t test; and (M) nose pokes in the last five minutes, *p=0.0116, two-tailed unpaired t test. *Cntnap2^+/+^* n=25 mice, *Cntnap2^−/−^* n=22 mice. (N) Quantification (mean ± SEM) of the frequency of movement syllables (top 25 most frequent syllables) in the open field assay defined by Keypoint-MoSeq. *Cntnap2^+/+^* n=13 mice and *Cntnap2^−/−^* n=11 mice, **p=0.0013 for grooming 6, **p=0.0013 for grooming 7, Kruskal-Wallis test with Dunn’s correction for multiple comparisons. (O) Quantification (mean ± SEM) of the entropy of syllable transitions in the open field assay. *Cntnap2^+/+^* n=13 mice and *Cntnap2^−/−^* n=11 mice, *p=0.0236, two-tailed unpaired t test. (P) Accuracy of a Random Forest decoder trained on DeepLabCut basic locomotor data (Movement), Keypoint-MoSeq syllable usage data (Syllable usage), or Keypoint-MoSeq syllable transition data (Syllable transition) in distinguishing between *Cntnap2^+/+^* and *Cntnap2^−/−^* mice. Dotted line represents chance performance. For panels E-M and O, dots/squares represent the value for each mouse.

To further assess motor behaviors in *Cntnap2^−/−^* mice, we utilized the marble burying assay (Fig. 5C), which takes advantage of a mouse’s natural tendency to dig or bury. The number of marbles buried is used as a measure of persistent or repetitive behavior (Angoa-Perez et al., 2013). We found that *Cntnap2^−/−^* mice buried significantly more marbles on average than WT controls (Fig. 5J). Another measure of repetitive behavior, which is based on the natural exploratory behavior of mice is the holeboard assay (Fig. 5D). In this test, the number of nose pokes made into unbaited holes is recorded. *Cntnap2^−/−^* mice made significantly more nose pokes within a 10-minute period than WT mice (Fig. 5K). This was largely due to increased poking during the last 5 minutes of the test (Fig. 5L-M), indicating persistent poking behavior in *Cntnap2^−/−^* mice. Together, the increased grooming, marble burying and nose poking indicate an increase in RRBs in *Cntnap2^−/−^* mice. A summary of behavior test results by genotype and sex is shown in Table S1.

To gain further insight into the spontaneous behavior profile of *Cntnap2^−/−^* mice, we utilized a combination of DeepLabCut and Keypoint-MoSeq to perform unbiased, machine learning-based assessment of general locomotion and behavior in an additional cohort of *Cntnap2^−/−^* mice (Fig. 5N-P, Fig. S7) (Mathis et al., 2018; Wiltschko et al., 2020). Again, we found that *Cntnap2^−/−^* mice did not exhibit changes in general locomotor activity compared to WT littermates (Fig. S7A). Analysis of movement “syllables” using Keypoint-MoSeq revealed that across the 25 most frequent syllables, two syllables associated with grooming were performed with significantly increased frequency in *Cntnap2^−/−^* mice (Fig. 5N). *Cntnap2^−/−^* mice also had an increase in the total number of grooming bouts (Fig. S7B), replicating the findings in the manually scored cohort (see Fig. 5I). While syllable usage was generally similar between WT and *Cntnap2^−/−^* mice, transitions between syllables differed between the groups (Fig. S7C). A measure of the entropy of syllable transitions revealed that *Cntnap2^−/−^* mice exhibited less entropy, suggesting less variability in the transition from one movement syllable to the next (Fig. 5O). This rigidity in motor sequence may be indicative of more restricted motor behavior overall. Finally, we tested whether a trained decoder could accurately distinguish WT and *Cntnap2^−/−^* mice using information about movement, syllable usage, or syllable transitions. The decoding models performed significantly better than chance at identifying WT and *Cntnap2^−/−^* mice based on their syllable usage and transitions, but not general locomotor activity (Fig. 5P). Together, this analysis demonstrates that while overall locomotor activity is not strongly affected in *Cntnap2^−/−^* mice, the behavior patterns of these mice are distinct from WT, reflecting enhanced repetitive behaviors.

### *Cntnap2^−/−^* mice exhibit enhanced motor learning

The accelerating rotarod is a striatal-dependent measure of motor coordination and learning that has been used across a range of ASD mouse models (Cording & Bateup, 2023). Changes in corticostriatal circuits have been identified in mouse models of ASD with altered performance in the task (Cording & Bateup, 2023). Given the altered corticostriatal drive in *Cntnap2^−/−^* mice, we tested whether motor coordination and learning were affected in these mice. In the rotarod test, mice learn to walk and then run to stay on a rotating rod as it increases in speed over the course of five minutes. Mice perform three trials a day for four days. In trials one through six, the rod increases in speed from five to 40 revolutions per minute (RPM), while in trials seven through 12 the rod increases from 10 to 80 RPM (Fig. 6A). Learning occurs over trials within a day, as well as across days, as the mouse develops and hones a stereotyped motor pattern to stay on the rod for increasing amounts of time (Rothwell et al., 2014; Yin et al., 2009). We found that *Cntnap2^−/−^* mice performed significantly better than WT mice in this task, particularly in the later trials when the rod rotates at the faster 10 to 80 RPM speed (Fig. 6B).

**Figure 6.**
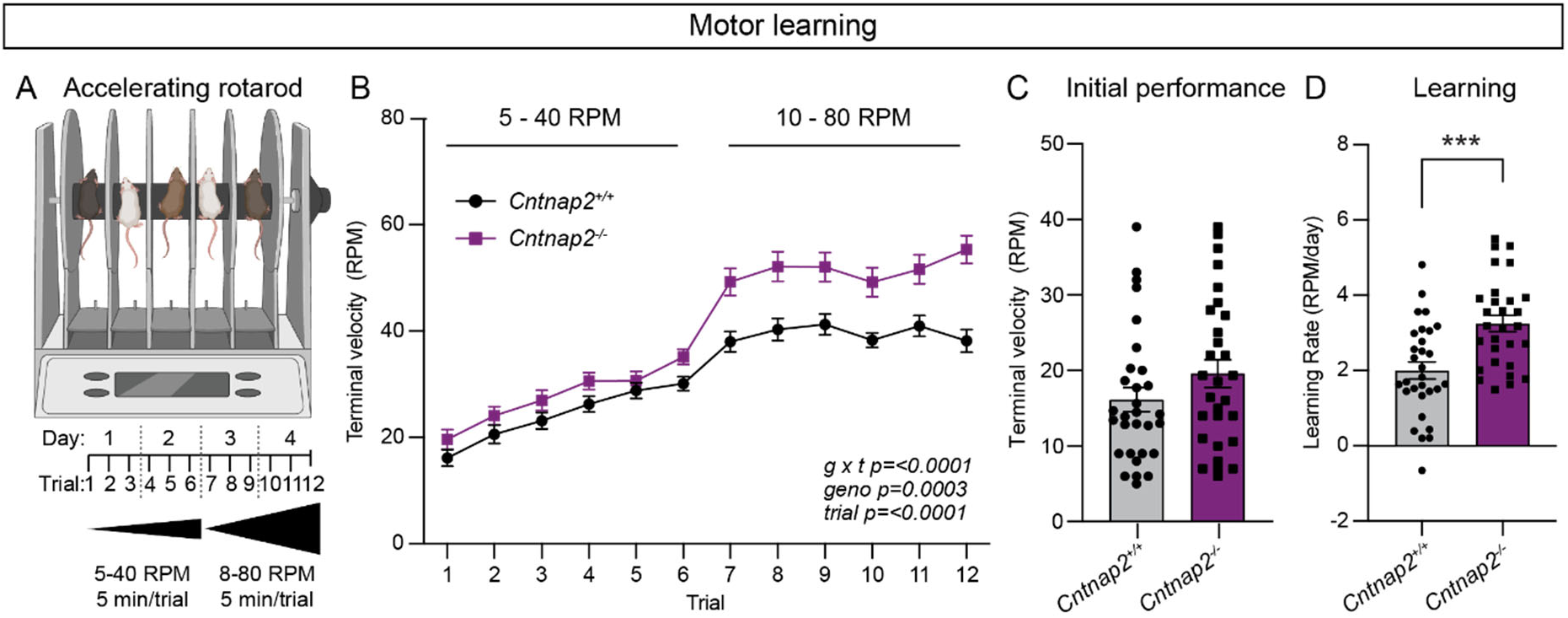
*Cntnap2^−/−^* mice exhibit enhanced motor learning. (A) Schematic of the rotarod apparatus (top), and design of the task (bottom), created with BioRender.com. Mice walk to stay on the rotating rod for three 5-minute trials a day for two days at 5-40 RPM acceleration over 5 minutes, followed by three trials a day for two days at 10-80 RPM. (B) Quantification (mean ± SEM) of accelerating rotarod performance across 12 trials for the indicated genotypes. *Cntnap2^+/+^* n=30 mice, *Cntnap2^−/−^* n=29 mice. Repeated measures two-way ANOVA p values are shown; g x t F (11, 616) = 4.935, geno F (1, 56) = 15.29, trial F (7.245, 405.7) = 108.4. (C) Quantification (mean ± SEM) of rotarod performance on trial 1 quantified as terminal speed. Dots/squares represent the performance of individual mice. n is same as in panel B, p=0.1518, Mann-Whitney test. (D) Quantification (mean ± SEM) of learning rate (RPM/day) calculated as the slope of the line of performance from the first trial (1) to the last trial (12) for each mouse. Dots/squares represent the learning rate for individual mice. n is same as in panel B, ***p=0.0002, two-tailed unpaired t test.

Initial performance (terminal velocity on trial one) was not significantly different between WT and *Cntnap2^−/−^* mice (Fig. 6C), but the learning rate from trial one to trial 12 was increased in *Cntnap2^−/−^* mice (Fig. 6D). These findings expand upon previous work indicating increased performance on both steady-state and accelerating rotarod tasks utilizing slower speeds in *Cntnap2^−/−^* mice (Dawes et al., 2018; Penagarikano et al., 2011). These results also align with the increased rotarod performance seen in other ASD mouse models exhibiting enhanced corticostriatal drive (Benthall et al., 2021; Cording & Bateup, 2023).

### *Cntnap2^−/−^* mice exhibit cognitive inflexibility

RRBs include not just stereotyped movements, but also insistence on sameness and perseverative interests (APA, 2022). Cognitive inflexibility, a deficit in the ability to flexibly adapt to changes in the environment and update behavior, is a manifestation of ASD and some other psychiatric disorders, which are associated with striatal dysfunction (Fuccillo, 2016). Indeed, in individuals with ASD, the severity of RRBs is associated with measures of cognitive inflexibility, and evidence from imaging studies suggests that altered corticostriatal connectivity may be present in the case of both repetitive behaviors and cognitive inflexibility (Uddin, 2021). To assess cognitive flexibility in *Cntnap2^−/−^* mice, we utilized a four-choice odor-based reversal learning assay (Johnson et al., 2016; Lin et al., 2022). Briefly, mice were trained to dig for a reward in one of four pots containing scented wood shavings (Fig. 7A). On the first day of the task (acquisition), the rewarded pot was scented with odor one (O1). Mice reached criterion when they chose O1 for at least eight of 10 consecutive trials. On day two, mice were given a recall test in which the rewarded pot was again scented with O1. After reaching criterion, the reversal trials began, and the rewarded pot was scented with the previously unrewarded odor two (O2). To reach criterion, mice must learn the new association of O2 and reward and choose O2 for eight of 10 consecutive trials.

**Figure 7.**
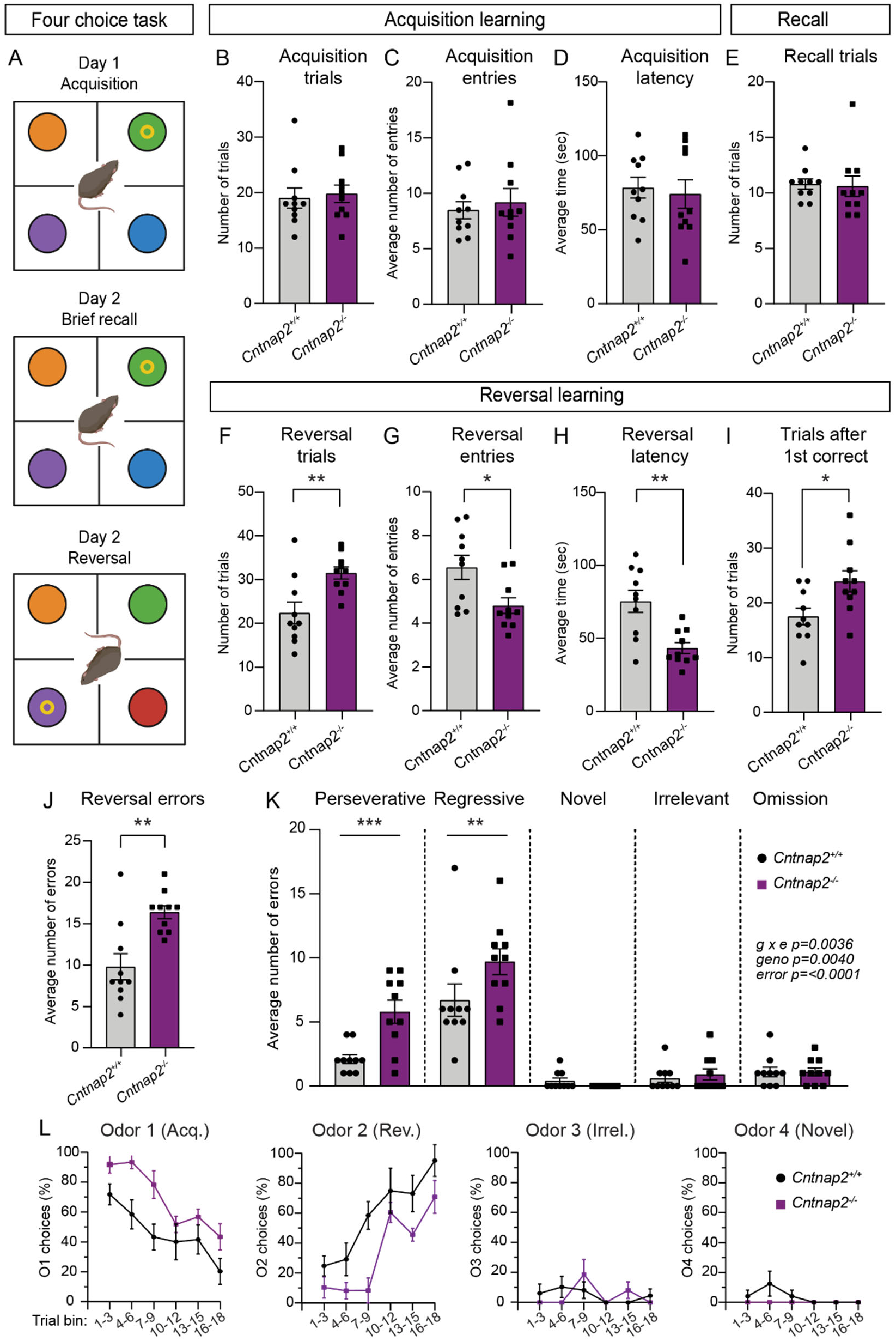
*Cntnap2^−/−^* mice demonstrate cognitive inflexibility. (A) Schematic of the four-choice odor-based reversal learning task, created with BioRender.com. Colored circles represent pots with different scented wood shavings. Yellow ring represents the food reward. Red circle in the Day 2 Reversal panel indicates a novel odor. (B-D) Quantification of parameters during acquisition learning. Mean ± SEM number of trials to reach criterion (B, at least 8 out of last 10 trials correct), p=0.5397, Mann-Whitney test; number of quadrant entries before making a choice (C), p=0.9118, Mann-Whitney test; and latency to make a choice (D), p=0.7224, two-tailed unpaired t test. Dots/squares represent the value for each mouse. (E) Quantification (mean ± SEM) of the number of trials to reach criterion (at least 8 out of last 10 trials correct) during the recall test on day 2, p=0.3737, Mann-Whitney test. (F-I) Quantification of parameters during reversal learning. Mean ± SEM number of trials to reach criterion (F, at least 8 out of last 10 trials correct), **p=0.0048, two-tailed unpaired t test; number of quadrant entries before making a choice (G), *p=0.0158, two-tailed unpaired t test; latency to make a choice (H), **p=0.0013, two-tailed unpaired t test; and number of trials to reach criterion after the first correct choice (I), *p=0.0183, two-tailed unpaired t test. (J) Quantification (mean ± SEM) of the total number of errors made during reversal learning, **p=0.0034, Mann-Whitney test. (K) Quantification (mean ± SEM) of the different error types made during reversal learning. Perseverative errors, ***p=0.0005, regressive errors **p=0.0068, novel errors, p=0.9955, irrelevant errors p=0.9988, omissions, p=>0.9999, repeated measures two-way ANOVA with Šídák’s multiple comparisons test; g x e F (4, 72) = 4.292, geno F (1, 18) = 10.91, error F (4, 72) = 53.49. (L) Quantification (mean ± SEM) of the percent of choices made for each odor, binned across three trials, during reversal learning. Odor 1 was rewarded during acquisition learning. Odor 2 was rewarded during reversal learning. Odor 3 was never rewarded (irrelevant). Odor 4 was a novel odor introduced during the reversal learning phase. For panels B-L, n=10 *Cntnap2^+/+^* mice and 10 *Cntnap2^−/−^* mice.

During acquisition, *Cntnap2^−/−^* mice performed similarly to WT controls, not differing in the average number of trials needed to reach criterion, the number of quadrant entries made before making a choice, or the latency to choose a pot (Fig. 7B-D). On day two, *Cntnap2^−/−^* mice performed similarly to controls during recall, demonstrating successful consolidation of the odor-reward pairing (Fig. 7E). However, we found that *Cntnap2^−/−^* mice had a deficit in reversal learning, requiring significantly more trials on average than WTs to reach criterion once the odor-reward pairing was changed (Fig. 7F). Interestingly, during reversal, *Cntnap2^−/−^* mice made fewer quadrant entries before making a digging choice and had significantly decreased latency to make a choice compared to controls (Fig. 7G,H). Even after the first correct choice of O2 during reversal, *Cntnap2^−/−^* mice took more trials to reach criterion than WTs (Fig. 7I). In terms of errors, *Cntnap2^−/−^* mice made more reversal errors than WT mice (Fig. 7J), in particular perseverative (continuing to choose O1) and regressive (choosing O1 after correctly choosing O2 once) errors (Fig. 7K). *Cntnap2^−/−^* mice did not differ from WT controls in choices of the novel (newly introduced during reversal) or irrelevant (never rewarded) odors, or in the number of omitted trials (timing out without making a choice) (Fig. 7K). Instead, the persistence in choosing O1, even after at least one correct choice of O2, drove the cognitive inflexibility in these mice (Fig. 7L). This persistence in choice may be reflective of the broader scope of RRBs in *Cntnap2^−/−^* mice.

## Discussion

In this study we tested whether loss of the ASD risk gene *Cntnap2* altered striatal physiology or striatal-dependent behaviors. We found that direct pathway SPNs exhibited enhanced cortical drive in *Cntnap2^−/−^* mice. This change was not due to differences in excitatory or inhibitory synapses, as cortical inputs onto SPNs were unchanged and there were no significant deficits in inhibition onto SPNs in these mice. Instead, loss of *Cntnap2* resulted in a significant increase in the excitability of dSPNs, likely driven by altered contribution of Kv1.2 channels to intrinsic firing properties. At the behavioral level, *Cntnap2^−/−^* mice exhibited repetitive behaviors including increased grooming, nose poking, and marble burying. These mice also had enhanced motor learning, performing significantly better than controls in the accelerating rotarod task. Finally, *Cntnap2^−/−^* mice exhibited cognitive inflexibility in the four-choice reversal learning assay.

### Cellular phenotypes of *Cntnap2* loss

The loss of Caspr2 has a variable impact on intrinsic excitability across brain regions and cell types. The increased intrinsic excitability that we identified in dSPNs has also been observed in Purkinje cells of the cerebellum (Fernandez et al., 2021) and pyramidal cells of the cortex (Antoine et al., 2019; Cifuentes-Diaz et al., 2023). However, we note hypoactivity (Brumback et al., 2018) or unchanged excitability (Lazaro et al., 2019) of pyramidal cells in some cortical regions in *Cntnap2^−/−^* mice. Caspr2 is involved in the clustering of voltage-gated potassium channels, in particular at the juxtaparanodes of myelinated axons (Poliak et al., 1999; Poliak et al., 2003) and the axon initial segment (Inda et al., 2006). Indeed, there are profound deficits in the clustering of Kv1-family channels in *Cntnap2^−/−^* mice, particularly Kv1.2 channels (Scott et al., 2019). These channels play an important role in regulating the intrinsic excitability of SPNs (Nisenbaum et al., 1994), in particular dSPNs (Lahiri & Bevan, 2020), and when blocked, result in increased excitability (Shen et al., 2004). The loss of Caspr2 in *Cntnap2^−/−^* mice may result in improper localization of Kv1.2 channels, and thus alter their contribution to intrinsic excitability. Indeed, we found that the effects of Kv1.2 blockade were largely occluded in *Cntnap2^−/−^* dSPNs, indicating that Kv1.2 loss of function is likely the mechanism driving the change in excitability. Interestingly, we find that blocking Kv1.2 channels has less of an effect on the excitability of iSPNs, which may account for the greater impact of *Cntnap2* loss on dSPNs.

Prior studies of *Cntnap2^−/−^* mice have identified changes in the number of PV-expressing interneurons in the cortex (Penagarikano et al., 2011; Vogt et al., 2018), hippocampus (Paterno et al., 2021; Penagarikano et al., 2011) and striatum (Penagarikano et al., 2011). However, this finding is inconsistent across studies, as others have reported no change in the number of PV interneurons in these regions (Ahmed et al., 2023; Lauber et al., 2018; Scott et al., 2019). One possible explanation for this disparity is altered PV protein expression in *Cntnap2^−/−^* mice such that immunoreactivity varies in cell counting assessments. This is supported by the finding that the number of Vicia Villosa Agglutinin-positive (VVA+) perineuronal nets that preferentially surround PV cells is unchanged in *Cntnap2^−/−^* mice, even when PV immunoreactivity varies (Hartig et al., 1992; Haunso et al., 2000; Lauber et al., 2018). Parvalbumin, a Ca^2+^ buffer, plays an important role in the intrinsic fast-spiking properties of PV interneurons, such that a reduction in PV protein expression is known to change PV intrinsic function (Orduz et al., 2013). However, altered intrinsic properties of PV interneurons has also been variably reported across brain regions and studies of *Cntnap2^−/−^* mice, with subtle changes in PV firing properties reported in the developing striatum (Ahmed et al., 2023) and adult cortex (Vogt et al., 2018), but unchanged in the hippocampus (Paterno et al., 2021) and medial prefrontal cortex (Lazaro et al., 2019). In this study, we find no significant change in the number of PV interneurons or the striatal expression of PV protein in *Cntnap2^−/−^* mice. Consistent with this, we find no deficits in PV-mediated inhibition onto SPNs. Together, this suggests that primary changes in PV interneurons are unlikely to account for altered striatal circuit function in *Cntnap2^−/−^* mice.

### Loss of *Cntnap2* alters striatal-dependent behaviors

The striatum can be separated into functionally distinct subregions. We focused on the dorsal striatum in this study because of its role in controlling motor and cognitive functions (Voorn et al., 2004), which are relevant to ASD (Fuccillo, 2016; Subramanian et al., 2017). The dorsal striatum can be further subdivided into the dorsomedial striatum (DMS) and the dorsolateral striatum (DLS), with the former considered an associative region involved in goal-directed action-outcome learning and the latter implicated in the acquisition of habitual or procedural behaviors (Packard & Knowlton, 2002). We focused on cellular properties in the DLS as stereotyped, perseverative or persistent behaviors likely recruit DLS circuitry (Evans et al., 2024; Fuccillo, 2016). In the accelerating rotarod assay, learning and performance in the task has been associated with changes in the DLS. Positive modulation of the firing rate of DLS neurons occurs during rotarod training, in particular in later trials of the task, and synaptic potentiation of DLS SPNs in late training is necessary for intact performance (Yin et al., 2009). In line with this, lesions of the DLS impair both early and late rotarod learning (Yin et al., 2009). We found that *Cntnap2^−/−^* mice had increased rotarod performance, most notably at the later stages when DLS function is strongly implicated. Functionally, we also found increased cortical drive of DLS dSPNs in these mice, a change that was sufficient to increase rotarod performance in another mouse model with disruption of an ASD-risk gene (Benthall et al., 2021). Together, this supports a connection between the change observed in DLS SPN physiology and the increased motor routine learning in *Cntnap2^−/−^* mice, although this idea remains to be causally tested.

In terms of restricted, repetitive behaviors, we replicated prior studies showing increased spontaneous grooming in *Cntnap2^−/−^* mice (Penagarikano et al., 2011). Early evidence implicates the striatum in the control of the syntax or sequence of movements in a rodent grooming bout, such that very small lesions of DLS are capable of disrupting grooming (Cromwell & Berridge, 1996). However, recent work has also outlined roles for cellular modulation in DMS and ventral striatal Islands of Calleja in the control of grooming behavior (Ramirez-Armenta et al., 2022; Zhang et al., 2021). *Cntnap2^−/−^* mice also exhibited increased marble burying and nose poking.

The precise neurobiological substrates of these behaviors are yet unclear, but evidence linking increases in these behaviors to changes in cortico-striatal and amygdala-striatal function supports the notion that these behaviors may fit into a broader basal ganglia-associated RRB-like domain (Albelda & Joel, 2012; Lee et al., 2024).

In the four-choice reversal learning task, *Cntnap2^−/−^* mice showed no differences during the acquisition phase, suggesting that there were no broad deficits in learning. However, in the reversal stage of the task, *Cntnap2^−/−^* mice took significantly more trials to learn a new odor-reward pairing, owing primarily to continued choice of the previously rewarded odor. The DMS and ventral striatum (nucleus accumbens) have been shown to play an important role in reversal learning (Izquierdo et al., 2017), and in the four choice task specifically (Delevich et al., 2022). Additionally, decreased dopamine release in the DLS is associated with deficits in reversal learning in this task (Kosillo et al., 2019; Lin et al., 2022). Together, the learning phenotypes seen in *Cntnap2^−/−^* mice in the accelerating rotarod and reversal learning assay share an underlying rigidity in behavioral choice. In both cases, changes in striatal circuits likely underlie the repetitive, stereotyped behaviors.

In summary, our results fit into a model whereby divergent cellular changes in the striatum driven by a functionally diverse set of ASD risk genes similarly enhance corticostriatal drive, in particular, of the direct pathway. This in turn may facilitate striatal-dependent motor routine learning and behavioral perseveration. We speculate that a shared gain-of-function in striatal circuits may play a role in the formation of perseverative or repetitive behaviors in a subset of ASDs more broadly.

### Limitations and future directions

This study characterized multiple striatal cell types and synapses as well as striatum-associated behaviors in *Cntnap2^−/−^* mice for the first time. However, there remain open questions as to the impact that *Cntnap2* loss has on striatal function. Although we did not identify excitatory or inhibitory synaptic changes in *Cntnap2^−/−^* SPNs in this study, we only focused on a subset of these connections. While cortical inputs are a major source of excitation onto SPNs, there are other excitatory inputs onto these cells, such as from thalamus, that were not assessed in this study (Ding et al., 2008; Doig et al., 2010; Gerfen & Surmeier, 2011). In addition, while all intrastriatal GABAergic interneurons would have been sampled in the electrical stimulation experiments in this study, it is possible that interrogation of individual interneuron subtypes would reveal changes in specific inhibitory connections. Finally, major modulators of SPN activity such as cholinergic interneurons and dopaminergic inputs were not assessed here. These inputs have been implicated in several ASD mouse models (Kosillo & Bateup, 2021; Paval & Miclutia, 2021; Rapanelli et al., 2017), and changes in striatal cholinergic interneuron function have been identified in young (P21) *Cntnap2^−/−^* mice, making further study of this circuit particularly cogent (Ahmed et al., 2023).

The major cellular phenotype we observed was enhanced intrinsic excitability of *Cntnap2^−/−^* dSPNs. Given that Kv1.2 channels are known to be organized in part by Caspr2 (Poliak et al., 1999; Poliak et al., 2003; Scott et al., 2019) and the fact that blockade of Kv1.2 did not affect the excitability of *Cntnap2^−/−^* dSPNs, we conclude that loss of *Cntnap2* leads to the improper clustering, number, or function of Kv1.2. However, more direct measurement of the number and/or localization of these channels through imaging, or the function of these channels through voltage clamp measurement of potassium currents, would bolster this conclusion. Further, assessing whether similar occlusion of the effects of blocking Kv1.2 channels occurs in the cortical and cerebellar cell types that have also been shown to be hyperexcitable in *Cntnap2^−/−^* mice would support a more holistic understanding of the impact of *Cntnap2* loss on neuronal function (Antoine et al., 2019; Cifuentes-Diaz et al., 2023; Fernandez et al., 2021). Considering that these cell types also exhibit α-DTX-sensitive currents that boost AP firing, it is possible that the mechanism by which *Cntnap2* loss increases excitability may be shared (Guan et al., 2007; Haghdoust et al., 2007; Khavandgar et al., 2005).

Finally, while we did identify changes in several striatum-associated behaviors in *Cntnap2^−/−^* mice, a causative relationship between the physiological and behavioral changes that we identified has not been established. Given that hyperexcitability of the SPNs of the movement-initiating direct pathway is the primary physiological change we identified, testing whether this change is necessary (i.e. by decreasing dSPN activity in *Cntnap2^−/−^* mice through cell type-specific expression of inward rectifying potassium channel Kir2.1) or sufficient (i.e. by increasing dSPN activity in WT mice using cell type-specific G_q_-coupled (hM2Dq) DREADD activation) to alter behavior could illuminate the relationship between striatal function and behavior in *Cntnap2^−/−^* mice.

## Supplementary Figures and Table

**Supplementary Figure 1.**
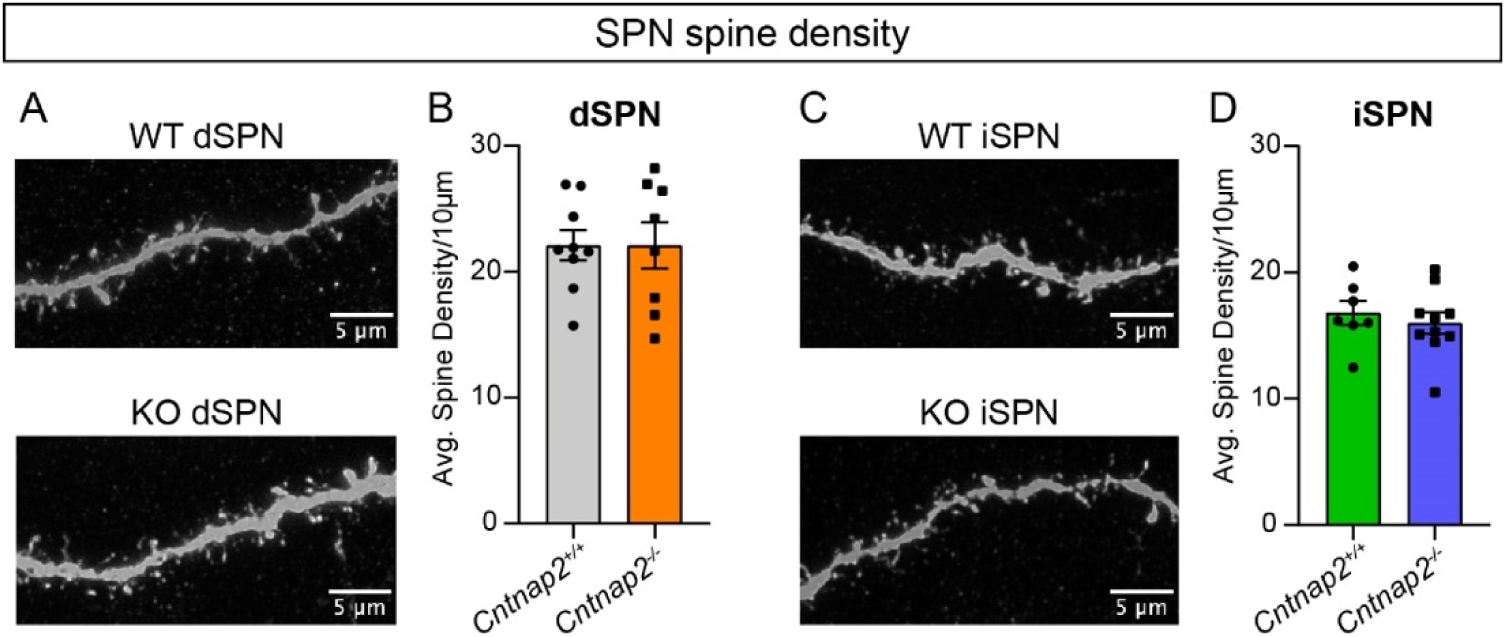
*Cntnap2^−/−^* SPNs do not have altered spine density. (A) Representative deconvoluted images of dendritic spines from dSPNs for the indicated genotypes. (B) Quantification (mean ± SEM) of dendritic spine density per 10 μm of dendrite in *Cntnap2^+/+^* and *Cntnap2^−/−^* dSPNs. Dots/squares represent the average spine density per neuron. *Cntnap2^+/+^* n=9 neurons (15 dendrites) from 6 mice, *Cntnap2^−/−^* n=8 neurons (15 dendrites) from 6 mice. p=0.9964, two-tailed unpaired t test. (C) Representative deconvoluted images of dendritic spines from iSPNs for the indicated genotypes. (D) Quantification (mean ± SEM) of dendritic spine density per 10 μm of dendrite in *Cntnap2^+/+^* and *Cntnap2^−/−^* iSPNs. Dots/squares represent the average spine density per neuron. *Cntnap2^+/+^* n=7 neurons (15 dendrites) from 4 mice, *Cntnap2^−/−^* n=10 neurons (15 dendrites) from 7 mice. p=0.5362, Mann-Whitney test.

**Supplementary Figure 2.**
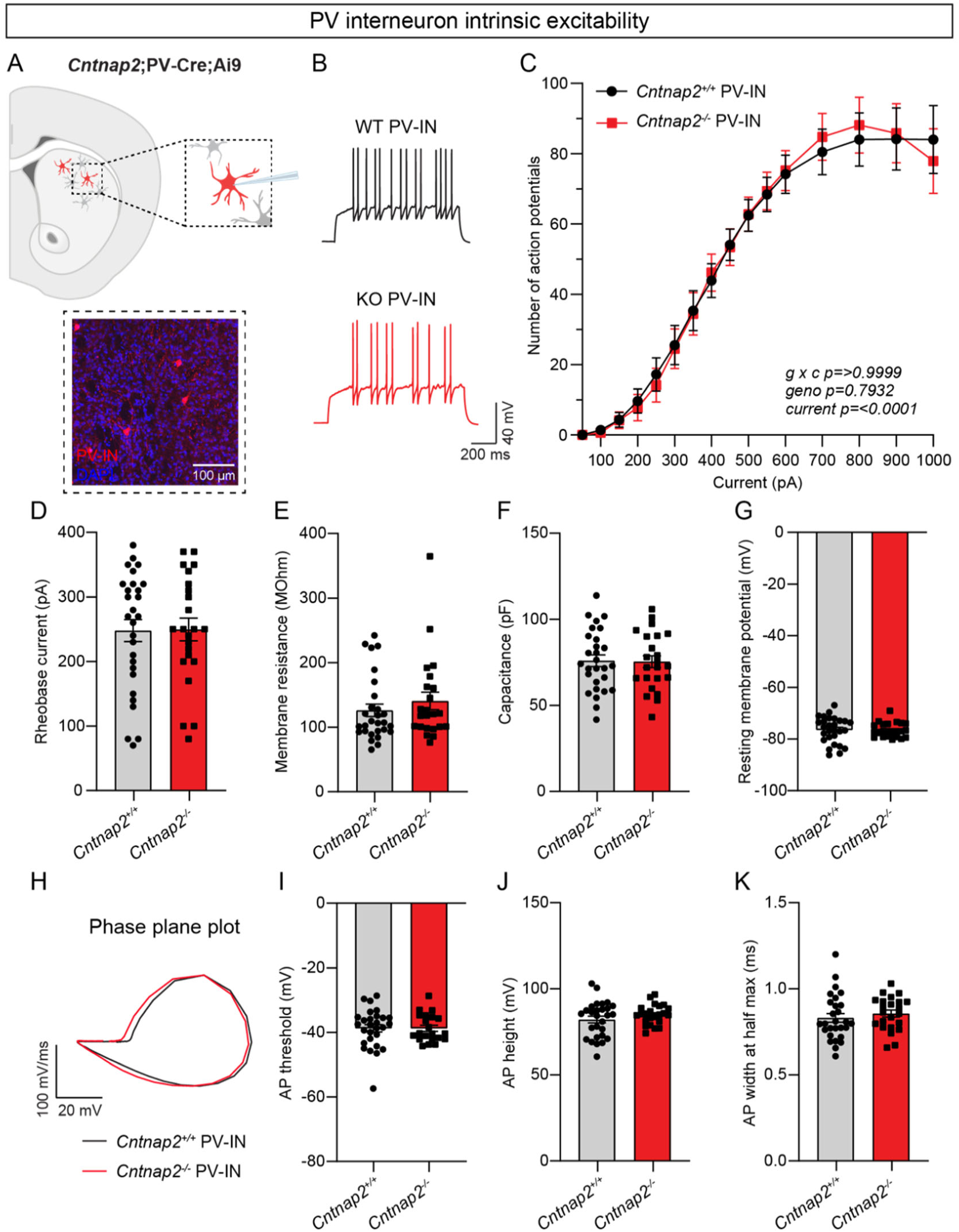
PV interneuron intrinsic excitability is unchanged in *Cntnap2^−/−^* mice. (A) Top: schematic of the experiment. tdTomato-expressing PV interneurons were recorded in the dorsolateral striatum and injected with current steps of increasing magnitude to evoke firing. Bottom: 20x confocal image of dorsolateral striatum from a *Cntnap2^+/+^;*PV-Cre;Ai9 mouse. tdTomato (red) labels PV interneurons, DAPI stained nuclei are in blue. (B) Example traces of APs in PV interneurons evoked by a 200 pA current step for the indicated genotypes. (C) Quantification (mean ± SEM) of the number of APs evoked in PV interneurons at different current step amplitudes. *Cntnap2^+/+^* n=28 cells from 6 mice, *Cntnap2^−/−^* n=23 cells from 4 mice. Repeated measures two-way ANOVA p values are shown; g x c F (40, 1960) = 0.1251, geno F (1, 49) = 0.06951, current F (2.033, 99.60) = 107.5. (D) Quantification (mean ± SEM) of the rheobase current. Dots/squares represent the rheobase current for each neuron. n is the same as in panel C. p=0.8852, two-tailed unpaired t test. (E-G) Quantification (mean ± SEM) of the membrane resistance (E), p=0.3422, Mann-Whitney test; membrane capacitance (F), p=0.9055, two-tailed unpaired t test; and resting membrane potential (G), p=0.3141, two-tailed unpaired t test. Dots/squares represent the average value for each neuron. n is the same as in panel C. (H) Phase plane plot of a single AP evoked in PV interneurons from *Cntnap2^+/+^* (black) and *Cntnap2^−/−^* (red) mice. (I-K) Quantification (mean ± SEM) of the AP threshold (I), p=0.6025, two-tailed unpaired t test; AP height (J), p=0.2850, two-tailed unpaired t test; and AP width at half max (K), p=0.2561, two-tailed unpaired t test. Dots/squares represent the value for the first spike evoked at rheobase for each neuron. n is the same as in panel C.

**Supplementary Figure 3.**
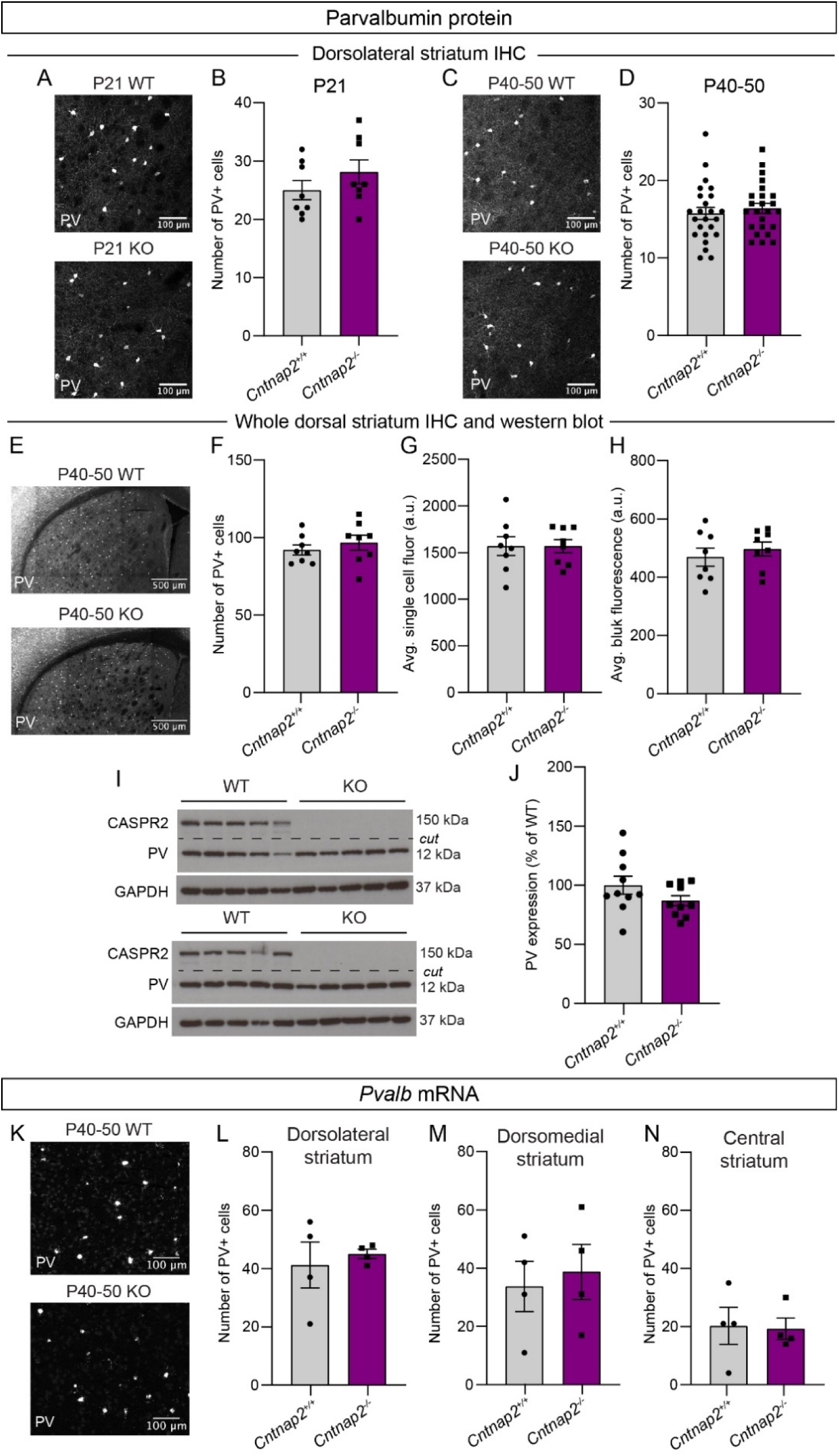
*Cntnap2^−/−^* mice do not exhibit changes in PV-positive cell number or expression. (A) Representative confocal images of dorsolateral striatum labeled with an antibody against Parvalbumin (PV) protein for the indicated genotypes, mice aged postnatal day (P) 21. (B) Quantification (mean ± SEM) of the number of PV positive cells per ROI at P21, p=0.2554, two-tailed unpaired t test. *Cntnap2^+/+^* n=8 sections from 4 mice (2 sections imaged per mouse), *Cntnap2^−/−^* n=8 sections from 4 mice (2 sections imaged per mouse). (C) Representative confocal images of dorsolateral striatum labeled with an antibody against PV for the indicated genotypes, mice aged P40-50. (D) Quantification (mean ± SEM) of the number of PV positive cells per ROI, p=0.5144, two-tailed unpaired t test. *Cntnap2^+/+^* n=24 sections from 12 mice (2 sections imaged per mouse), *Cntnap2^−/−^* n=24 sections from 12 mice (2 sections imaged per mouse). (E) Representative confocal images of dorsal striatum labeled with an antibody against PV for the indicated genotypes, mice aged P40-50. (F) Quantification (mean ± SEM) of the number of PV positive cells in the whole dorsal striatum at P40-50, p=0.4201, two-tailed unpaired t test. *Cntnap2^+/+^* n=8 sections from 4 mice (2 sections imaged per mouse), *Cntnap2^−/−^* n=8 sections from 4 mice (2 sections imaged per mouse). (G) Quantification (mean ± SEM) of the average single cell fluorescence of PV antibody staining in dorsal striatum, p=0.9996, two-tailed unpaired t test. *Cntnap2^+/+^* n=8 sections from 4 mice (2 sections imaged per mouse), *Cntnap2^−/−^* n=8 sections from 4 mice (2 sections imaged per mouse). (H) Quantification (mean ± SEM) of the average bulk fluorescence of PV antibody staining in dorsal striatum per section, p=0.4878, two-tailed unpaired t test. *Cntnap2^+/+^* n=8 sections from 4 mice (2 sections imaged per mouse), *Cntnap2^−/−^* n=8 sections from 4 mice (2 sections imaged per mouse). (I) Representative western blots for CASPR2, PV, and GAPDH protein in dorsal striatal tissue punches. 10 independent samples per genotype are shown. (J) Quantification (mean ± SEM) of PV protein expression normalized to GAPDH. Data are presented as a percentage of *Cntnap2^+/+^* levels, p=0.1485, two-tailed unpaired t test. *Cntnap2^+/+^* n=10 mice, *Cntnap2^−/−^* n=10 mice. (K) Representative confocal images of in situ hybridization for *Pvalb* in dorsolateral striatum for the indicated genotypes, mice aged P40-50. (L-N) Quantification (mean ± SEM) of the number of *Pvalb* positive cells in the dorsolateral striatum per section (L), p>0.9999, Mann-Whitney test; dorsomedial striatum (M), p=0.7429, Mann-Whitney test; and central striatum (N), p=0.6286, Mann-Whitney test. *Cntnap2^+/+^* n=4 sections from 2 mice (2 sections imaged per mouse), *Cntnap2^−/−^* n=4 sections from 2 mice (2 sections imaged per mouse).

**Supplementary Figure 4.**
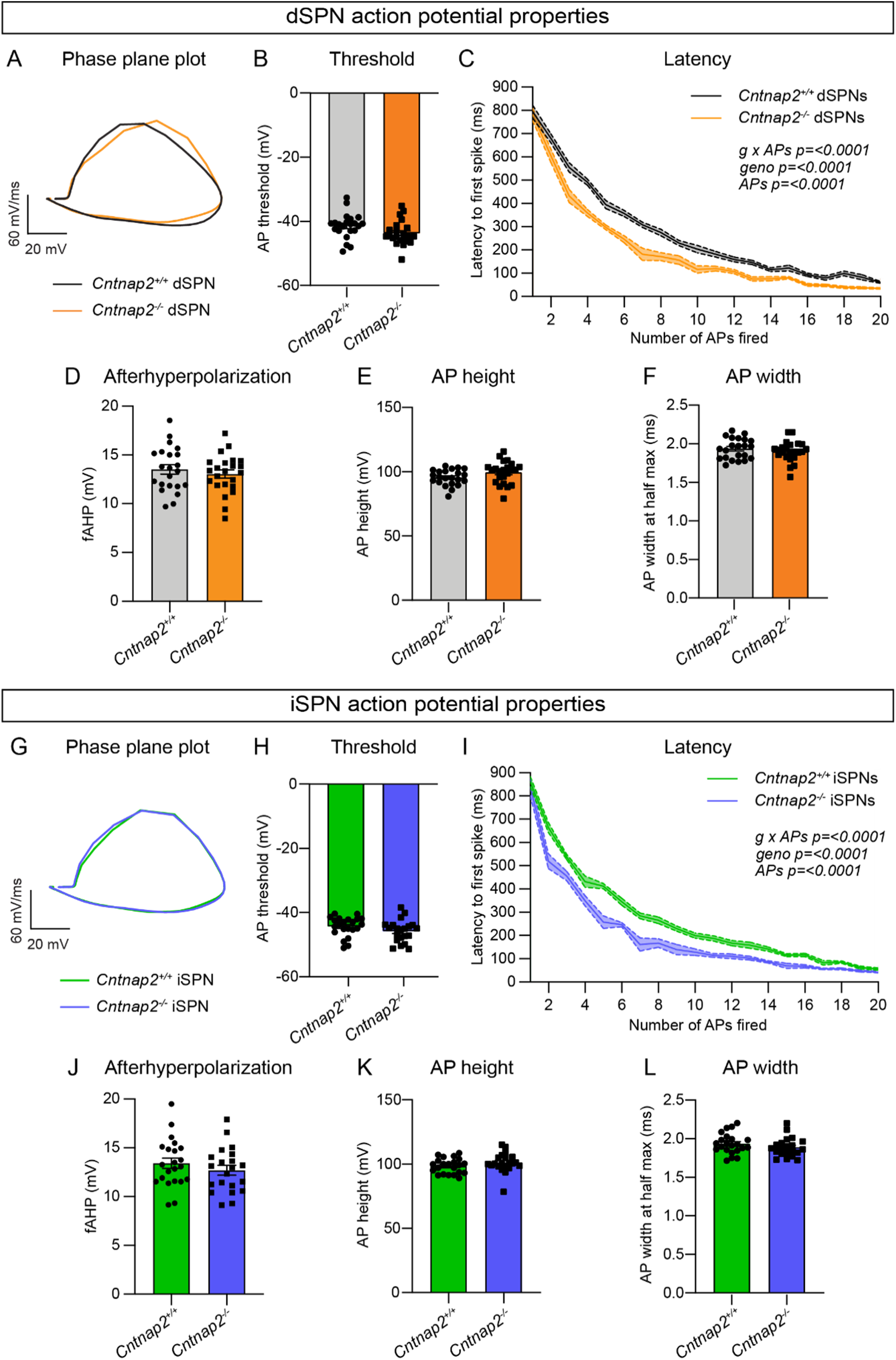
Latency to spike is reduced in *Cntnap2^−/−^* SPNs. (A) Phase plane plot of a single AP evoked in dSPNs from *Cntnap2^+/+^* (black) and *Cntnap2^−/−^* (orange) mice. (B) Quantification (mean ± SEM) of the dSPN AP threshold. Dots/squares represent the threshold for each neuron. *Cntnap2^+/+^* n=22 cells from 8 mice, *Cntnap2^−/−^* n=23 cells from 8 mice, p=0.0516, two-tailed unpaired t test. (C) Quantification (mean ± SEM) of dSPN AP latency to first spike plotted by number of APs fired in the current pulse. Shaded area around line represents SEM. Mixed-effects model with Geisser-Greenhouse correction p values are shown; g x APs F (19, 387) = 7.870, geno F (1, 43) = 24.98, APs F (1.740, 35.44) = 913.5. (D-F) Quantification (mean ± SEM) of the dSPN AP fast afterhyperpolarization (D), p=0.4664, two-tailed unpaired t test; AP height (E), p=0.0833, two-tailed unpaired t test; and AP width at half max (F), p=0.3792, two-tailed unpaired t test. Dots/squares represent the value for the first spike evoked at rheobase for each neuron, n same as in panel B. (G) Phase plane plot of a single AP evoked in iSPNs in *Cntnap2^+/+^* (green) and *Cntnap2^−/−^* (blue) mice. (H) Quantification (mean ± SEM) of the iSPN AP threshold. Dots/squares represent the threshold for each neuron. *Cntnap2^+/+^* n=22 cells from 8 mice, *Cntnap2^−/−^* n=21 cells from 8 mice, p=0.1495, two-tailed unpaired t test. (I) Quantification (mean ± SEM) of iSPN AP latency to first spike plotted by number of APs fired in the current pulse. Shaded area around line represents SEM. Mixed-effects model with Geisser-Greenhouse correction p values are shown; g x APs F (19, 343) = 3.300, geno F (1, 41) = 45.55, APs F (1.817, 32.81) = 680.4. (J-L) Quantification (mean ± SEM) of iSPN AP fast afterhyperpolarization (J), p=0.3281, two-tailed unpaired t test; AP height (K), p=0.1794, Mann-Whitney test; and AP width at half max (L), p=0.2409, two-tailed unpaired t test. Dots/squares represent the value for the first spike evoked at rheobase for each neuron, n same as in panel H.

**Supplementary Figure 5.**
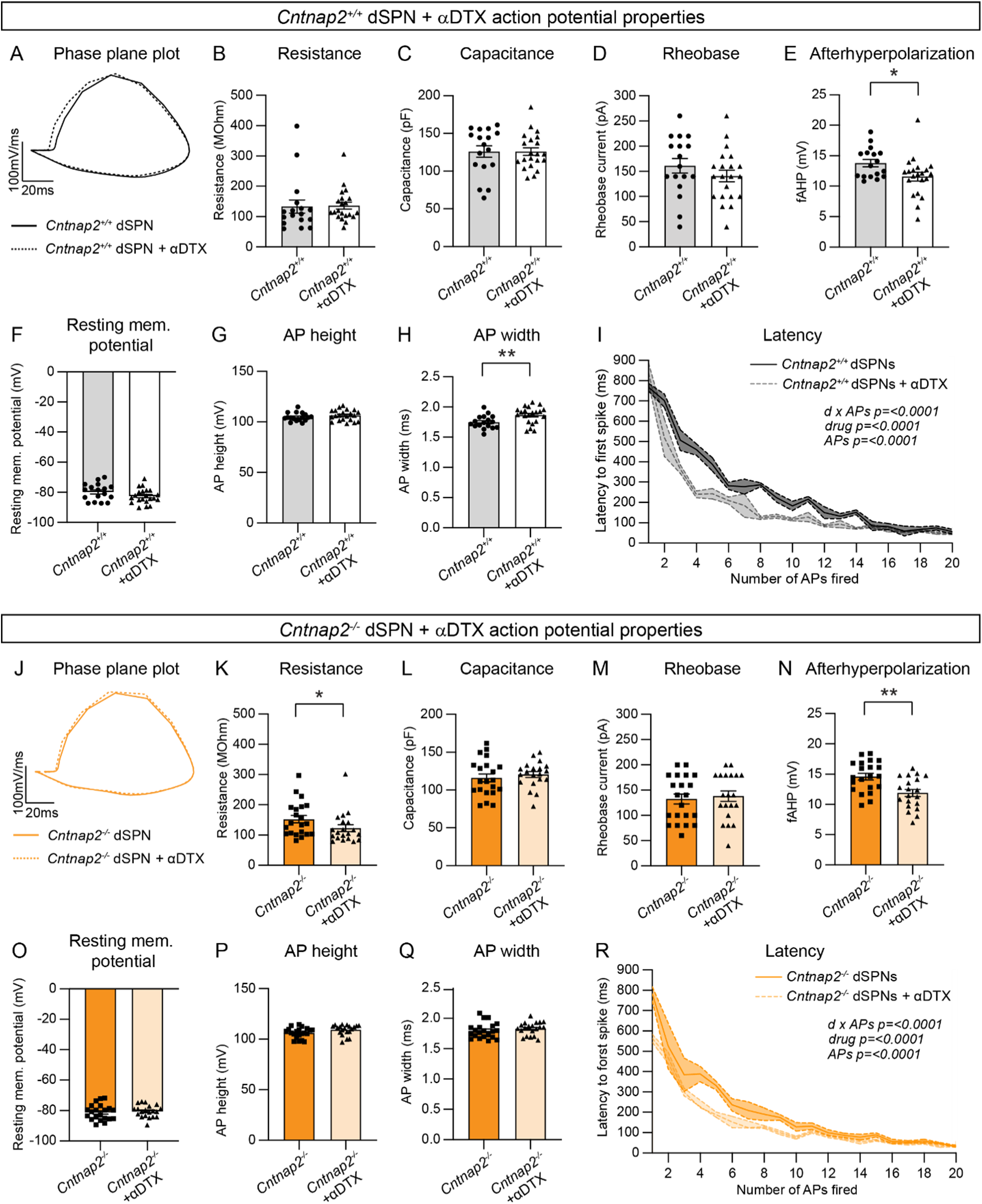
Inhibition of Kv1.2 differentially impacts action potential properties in *Cntnap2^+/+^* and *Cntnap2^−/−^* dSPNs. (A) Phase plane plot of a single AP evoked in *Cntnap2^+/+^* dSPNs in the absence (solid black) or presence (dotted black) of α-Dendrotoxin (α-DTX). (B-C) Quantification (mean ± SEM) of the *Cntnap2^+/+^* dSPN membrane resistance (B), p=0.2681, Mann-Whitney test; and membrane capacitance (C), p=0.9886, two-tailed unpaired t test. Dots/triangles represent the average value for each neuron, *Cntnap2^+/+^* n=17 cells from 8 mice, *Cntnap2^+/+^* + α-DTX n=21 cells from 8 mice. (D) Quantification (mean ± SEM) of the rheobase current in *Cntnap2^+/+^* dSPNs. Dots/triangles represent the rheobase current for each neuron. n is the same as in panels B-C. p=0.2703, two-tailed unpaired t test. (E-H) Quantification (mean ± SEM) of the *Cntnap2^+/+^* dSPN fast afterhyperpolarization (E), *p=0.0274, two-tailed unpaired t test; resting membrane potential (F), p=0.1207, two-tailed unpaired t test; AP height (G), p=0.3037, two-tailed unpaired t test; and AP width at half max (H), **p=0.0084, two-tailed unpaired t test. Dots/triangles represent the value for the first spike evoked at rheobase for each neuron. n is the same as in panels B-C. (I) Quantification (mean ± SEM) of *Cntnap2^+/+^* dSPN AP latency to first spike, plotted by number of APs fired in the current pulse. Shaded area around line represents SEM. Mixed-effects model with Geisser-Greenhouse correction p values are shown; d x APs F (19, 126) = 4.825, drug F (1, 36) = 23.33, APs F (19.00, 126.0) = 106.7. (J) Phase plane plot of a single AP evoked in *Cntnap2^−/−^* dSPNs in the absence (solid orange) or presence (dotted orange) of α-Dendrotoxin (α-DTX). (K-L) Quantification (mean ± SEM) of the *Cntnap2^−/−^* dSPNs membrane resistance (K), *p=0.0452, Mann-Whitney test; and membrane capacitance (L), p=0.4950, two-tailed unpaired t test. Squares/triangles represent the average value for each neuron, *Cntnap2^−/−^* n=21 cells from 9 mice, *Cntnap2^−/−^* + α-DTX n=20 cells from 9 mice. (M) Quantification (mean ± SEM) of the rheobase current in *Cntnap2^−/−^* dSPNs. Squares/triangles represent the rheobase current for each neuron. n is the same as in panels K-L. p=0.6239, Mann-Whitney test. (N-Q) Quantification (mean ± SEM) of the *Cntnap2^−/−^* dSPN fast afterhyperpolarization (N), **p=0.0017, two-tailed unpaired t test; resting membrane potential (O), p=0.5681, two-tailed unpaired t test; AP height (P), p=0.0652, two-tailed unpaired t test; and AP width at half max (Q), p=0.3400, two-tailed unpaired t test. Squares/triangles represent the value for the first spike evoked at rheobase for each neuron. n is the same as in panels K-L. (R) Quantification (mean ± SEM) of *Cntnap2^−/−^* dSPN AP latency to first spike, plotted by number of APs fired in the current pulse. Shaded area around line represents SEM. Mixed-effects model with Geisser-Greenhouse correction p values are shown; d x APs F (19, 135) = 3.362, drug F (1, 39) = 30.66, APs F (19.00, 135.0) = 123.8.

**Supplementary Figure 6.**
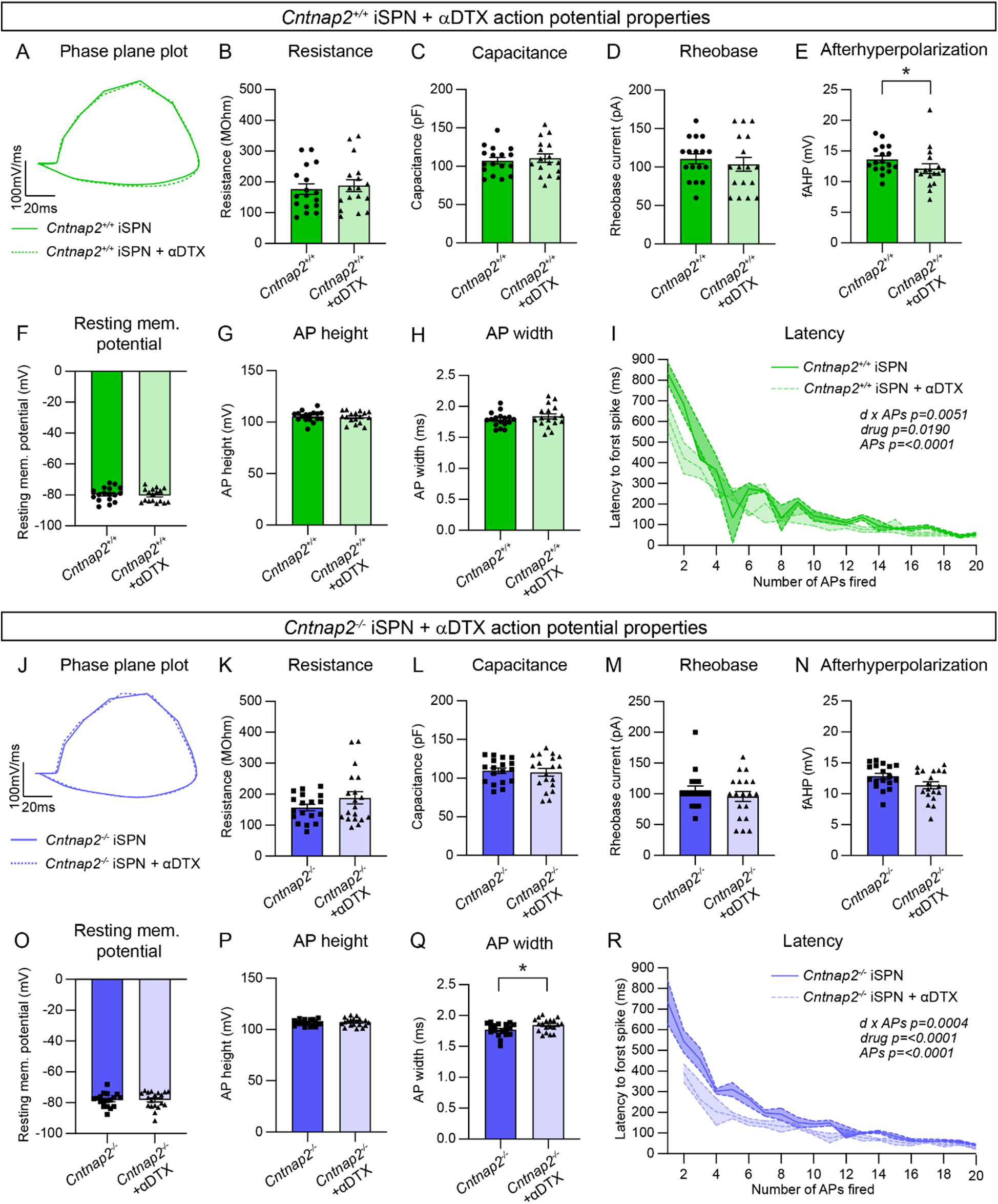
Inhibition of Kv1.2 does not strongly affect the excitability of iSPNs. (A) Phase plane plot of a single AP evoked in *Cntnap2^+/+^* iSPNs in the absence (solid green) or presence (dotted green) of α-Dendrotoxin (α-DTX). (B-C) Quantification (mean ± SEM) of the *Cntnap2^+/+^* iSPN membrane resistance (B), p=0.6624, two-tailed unpaired t test; and membrane capacitance (C), p=0.6444, two-tailed unpaired t test. Dots/triangles represent the average value for each neuron, *Cntnap2^+/+^* n=17 cells from 9 mice, *Cntnap2^+/+^* + α-DTX n=17 cells from 9 mice. (D) Quantification (mean ± SEM) of the rheobase current in *Cntnap2^+/+^* iSPNs. Dots/triangles represent the rheobase current for each neuron. n is the same as in panels B-C. p=0.5267, two-tailed unpaired t test. (E-H) Quantification (mean ± SEM) of the *Cntnap2^+/+^* iSPN fast afterhyperpolarization (E), *p=0.0238, Mann-Whitney test; resting membrane potential (F), p=0.6832, Mann-Whitney test; AP height (G), p=0.6448, two-tailed unpaired t test; and AP width at half max (H), p=0.3070, two-tailed unpaired t test. Dots/triangles represent the value for the first spike evoked at rheobase for each neuron. n is the same as in panels B-C. (I) Quantification (mean ± SEM) of *Cntnap2^+/+^* iSPN AP latency to first spike, plotted by number of APs fired in the current pulse. Shaded area around line represents SEM. Mixed-effects model with Geisser-Greenhouse correction p values are shown; d x APs F (17, 82) = 2.375, drug F (1, 32) = 6.104, APs F (17.00, 82.00) = 40.22. (J) Phase plane plot of a single AP evoked in *Cntnap2^−/−^* iSPNs in the absence (solid blue) or presence (dotted blue) of α-Dendrotoxin (α-DTX). (K-L) Quantification (mean ± SEM) of the *Cntnap2^−/−^* iSPNs membrane resistance (K), p=0.1746, two-tailed unpaired t test; and membrane capacitance (L), p=0.7511, two-tailed unpaired t test. Squares/triangles represent the average value for each neuron, *Cntnap2^−/−^* n=18 cells from 10 mice, *Cntnap2^−/−^* + α-DTX n=19 cells from 10 mice. (M) Quantification (mean ± SEM) of the rheobase current in *Cntnap2^−/−^* iSPNs. Squares/triangles represent the rheobase current for each neuron. n is the same as in panels K-L. p=0.6589, Mann-Whitney test. (N-Q) Quantification (mean ± SEM) of the *Cntnap2^−/−^* iSPN fast afterhyperpolarization (N), p=0.0538, two-tailed unpaired t test; resting membrane potential (O), p=0.9461, two-tailed unpaired t test; AP height (P), p=0.5920, two-tailed unpaired t test; and AP width at half max (Q), *p=0.0335, two-tailed unpaired t test, in the absence or presence of α-DTX. Squares/triangles represent the value for the first spike evoked at rheobase for each neuron. n is the same as in panels K-L. (R) Quantification (mean ± SEM) of *Cntnap2^−/−^* iSPN AP latency to first spike in the absence or presence of α-DTX, plotted by number of APs fired in the current pulse. Shaded area around line represents SEM. Mixed-effects model with Geisser-Greenhouse correction p values are shown; d x APs F (17, 68) = 3.181, drug F (1, 35) = 20.66, APs F (17.00, 68.00) = 82.73.

**Supplementary Figure 7.**
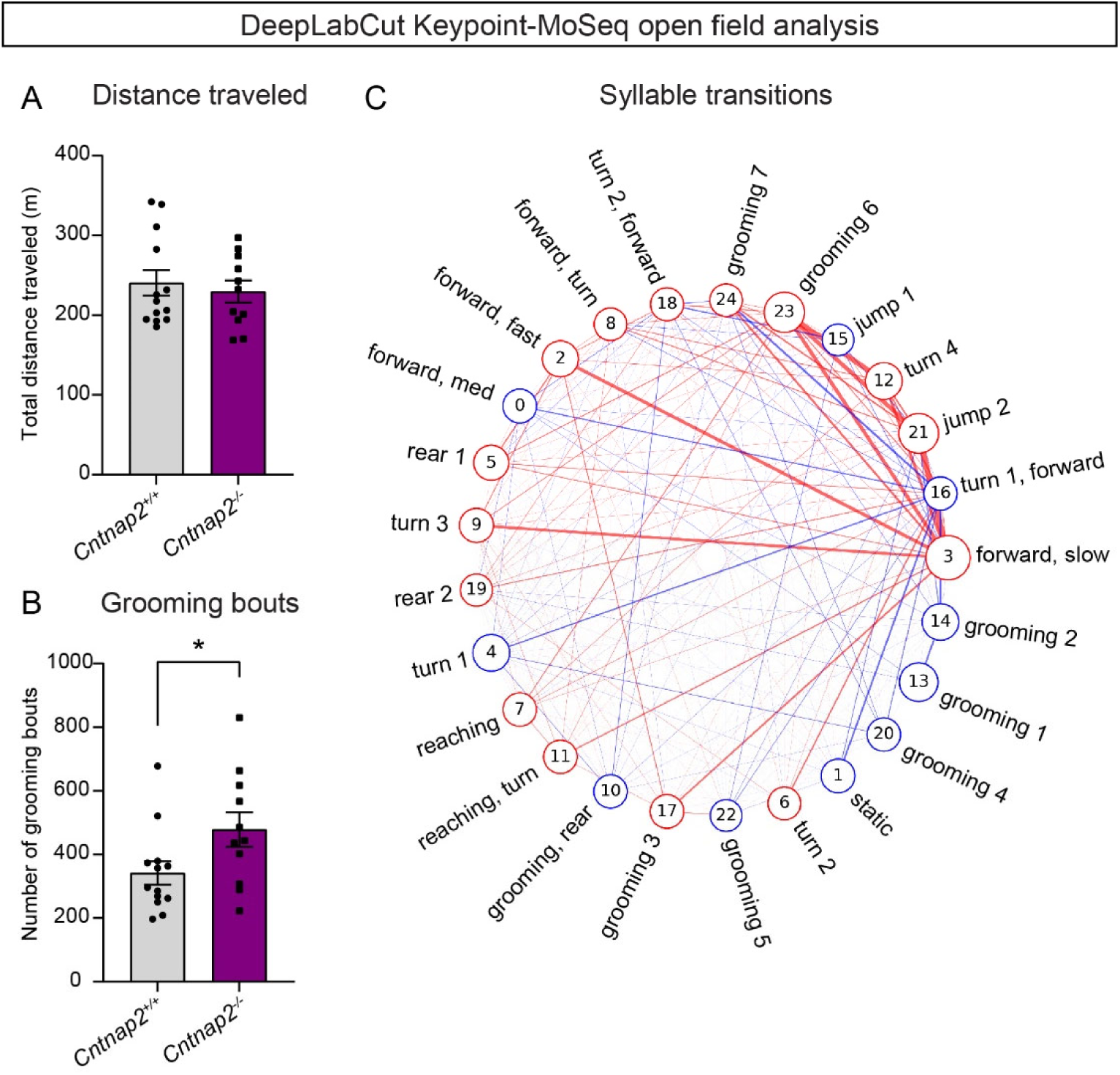
DeepLabCut and Keypoint-MoSeq analysis of *Cntnap2^−/−^* mice. (A) Quantification (mean ± SEM) of the total distance traveled in the open field measured by DeepLabCut, p=0.8201, Mann-Whitney test; *Cntnap2^+/+^* n=13 mice and *Cntnap2^−/−^* n=11 mice. (B) Quantification (mean ± SEM) of the number of grooming bouts in the open field assessed by Keypoint-MoSeq, calculated as the number of instances of all grooming syllables. *Cntnap2^+/+^* n=13 mice and *Cntnap2^−/−^* n =11 mice, *p = 0.0352, Mann-Whitney test. (C) Map depicting the difference in transitions between syllables (top 25 most frequent syllables) for *Cntnap2^+/+^* and *Cntnap2^−/−^* mice (represented as *Cntnap2^−/−^* minus *Cntnap2^+/+^*). A red circle around a syllable number indicates increased usage of that syllable in *Cntnap2^−/−^* mice, a blue circle indicates decreased usage of that syllable in *Cntnap2^−/−^* mice. A red line between syllable numbers indicates increased usage of that syllable transition in *Cntnap2^−/−^* mice, a blue line indicates decreased usage of that syllable transition in *Cntnap2^−/−^* mice. The thickness of the line scales with the size of the difference and the size of the circle surrounding the syllable number scales with usage of the syllable.

**Supplementary Table 1.** Summary of behavior data by sex and genotype.

| **Group** | **N**<br>**(mice)** | **Mean** | **Std.**<br>**deviation** | **Std. error of**<br>**the mean** |
| --- | --- | --- | --- | --- |
| ***Open field*** |  |  |  |  |
| *Cntnap2*<sup>+/+</sup> male distance traveled | 22 | 127.7 m | 54.34 | 11.58 |
| *Cntnap2*<sup>-/-</sup> male distance traveled | 16 | 156 m | 28.36 | 7.091 |
| *Cntnap2*<sup>+/+</sup> female distance traveled | 19 | 139.2 m | 48.71 | 11.17 |
| *Cntnap2*<sup>-/-</sup> female distance traveled | 18 | 131.7 m | 57.52 | 13.56 |
| *Cntnap2*<sup>+/+</sup> male avg. speed | 22 | 0.03536 m/s | 0.01500 | 0.003198 |
| *Cntnap2*<sup>-/-</sup> male avg. speed | 16 | 0.04331 m/s | 0.007964 | 0.001991 |
| *Cntnap2*<sup>+/+</sup> female avg. speed | 19 | 0.03868 m/s | 0.01341 | 0.003076 |
| *Cntnap2*<sup>-/-</sup> female avg. speed | 18 | 0.03644 m/s | 0.01590 | 0.003749 |
| *Cntnap2*<sup>+/+</sup> male rears | 22 | 557.6 rears | 201.8 | 43.03 |
| *Cntnap2*<sup>-/-</sup> male rears | 16 | 462.8 rears | 173.0 | 43.25 |
| *Cntnap2*<sup>+/+</sup> female rears | 19 | 361.2 rears | 188.3 | 43.19 |
| *Cntnap2*<sup>-/-</sup> female rears | 18 | 355.1 rears | 158.8 | 37.42 |
| *Cntnap2*<sup>+/+</sup> male center entries | 22 | 254.3 entries | 123.3 | 26.29 |
| *Cntnap2*<sup>-/-</sup> male center entries | 16 | 350.8 entries | 102.4 | 25.59 |
| *Cntnap2*<sup>+/+</sup> female center entries | 19 | 204.8 entries | 71.51 | 16.41 |
| *Cntnap2*<sup>-/-</sup> female center entries | 18 | 247.6 entries | 88.30 | 20.81 |
| *Cntnap2*<sup>+/+</sup> male number of grooming bouts (20 min.) | 22 | 14.91 bouts | 8.602 | 1.834 |
| *Cntnap2*<sup>-/-</sup> male number of grooming bouts (20 min.) | 16 | 19.56 bouts | 11.52 | 2.879 |
| *Cntnap2*<sup>+/+</sup> female number of grooming bouts (20 min.) | 19 | 13.37 bouts | 6.353 | 1.457 |
| *Cntnap2*<sup>-/-</sup> female number of grooming bouts (20 min.) | 18 | 28.44 bouts | 18.99 | 4.475 |
| **Marble burying** |  |  |  |  |
| *Cntnap2*<sup>+/+</sup> male marbles buried | 16 | 8.125 marbles | 6.206 | 1.552 |
| *Cntnap2*<sup>-/-</sup> male marbles buried | 17 | 10.82 marbles | 5.187 | 1.258 |
| *Cntnap2*<sup>+/+</sup> female marbles buried | 17 | 8.235 marbles | 5.641 | 1.368 |
| *Cntnap2*<sup>-/-</sup> female marbles buried | 16 | 11.25 marbles | 5.310 | 1.328 |
| **Holeboard** |  |  |  |  |
| *Cntnap2*<sup>+/+</sup> male 10 min. nose pokes | 11 | 156.5 pokes | 21.18 | 6.385 |
| *Cntnap2*<sup>-/-</sup> male 10 min. nose pokes | 11 | 192.3 pokes | 38.79 | 11.70 |
| *Cntnap2*<sup>+/+</sup> female 10 min. nose pokes | 14 | 175.9 pokes | 37.51 | 10.03 |
| *Cntnap2*<sup>-/-</sup> female 10 min. nose pokes | 11 | 180.5 pokes | 47.50 | 14.32 |
| *Cntnap2*<sup>+/+</sup> male first 5 min. nose pokes | 11 | 105.4 pokes | 16.26 | 4.901 |
| *Cntnap2*<sup>-/-</sup> male first 5 min. nose pokes | 11 | 114.5 pokes | 30.25 | 9.121 |
| *Cntnap2*<sup>+/+</sup> female first 5 min. nose pokes | 14 | 110.5 pokes | 22.93 | 6.128 |
| *Cntnap2*<sup>-/-</sup> female first 5 min. nose pokes | 11 | 112.3 pokes | 30.09 | 9.073 |
| *Cntnap2*<sup>+/+</sup> male last 5 min. nose pokes | 11 | 51.18 pokes | 10.09 | 3.042 |
| *Cntnap2*<sup>-/-</sup> male last 5 min. nose pokes | 11 | 77.73 pokes | 14.80 | 4.462 |
| *Cntnap2*<sup>+/+</sup> female last 5 min. nose pokes | 14 | 65.36 pokes | 20.82 | 5.565 |
| *Cntnap2*<sup>-/-</sup> female last 5 min. nose pokes | 11 | 68.27 pokes | 20.15 | 6.077 |
| **Rotarod** |  |  |  |  |
| *Cntnap2*<sup>+/+</sup> male learning rate | 15 | 1.839 RPM/day | 1.416 | 0.3655 |
| *Cntnap2*<sup>-/-</sup> male learning rate | 14 | 2.856 RPM/day | 1.166 | 0.3115 |
| *Cntnap2*<sup>+/+</sup> female learning rate | 15 | 2.162 RPM/day | 1.083 | 0.2798 |
| *Cntnap2*<sup>-/-</sup> female learning rate | 15 | 3.607 RPM/day | 1.103 | 0.2849 |
| **Four choice reversal learning** |  |  |  |  |
| *Cntnap2*<sup>+/+</sup> male acquisition trials | 5 | 20.80 trials | 6.834 | 3.056 |
| *Cntnap2*<sup>-/-</sup> male acquisition trials | 5 | 20 trials | 7.106 | 3.178 |
| *Cntnap2*<sup>+/+</sup> female acquisition trials | 5 | 17.20 trials | 4.550 | 2.035 |
| *Cntnap2*<sup>-/-</sup> female acquisition trials | 5 | 19.60 trials | 2.302 | 1.030 |
| *Cntnap2*<sup>+/+</sup> male recall trials | 5 | 11 trials | 1.871 | 0.8367 |
| *Cntnap2*<sup>-/-</sup> male recall trials | 5 | 11 trials | 4.243 | 1.897 |
| *Cntnap2*<sup>+/+</sup> female recall trials | 5 | 10.60 trials | 1.140 | 0.5099 |
| *Cntnap2*<sup>-/-</sup> female recall trials | 5 | 10.20 trials | 1.095 | 0.4899 |
| *Cntnap2*<sup>+/+</sup> male reversal trials | 5 | 18.40 trials | 1.817 | 0.8124 |
| *Cntnap2*<sup>-/-</sup> male reversal trials | 5 | 30.40 trials | 4.037 | 1.806 |
| *Cntnap2*<sup>+/+</sup> female reversal trials | 5 | 26.40 trials | 9.737 | 4.354 |
| *Cntnap2*<sup>-/-</sup> female reversal trials | 5 | 32.60 trials | 4.827 | 2.159 |
| *Cntnap2*<sup>+/+</sup> male reversal errors | 5 | 7.400 errors | 0.8944 | 0.4000 |
| *Cntnap2*<sup>-/-</sup> male reversal errors | 5 | 17.60 errors | 2.608 | 1.166 |
| *Cntnap2*<sup>+/+</sup> female reversal errors | 5 | 12.20 errors | 6.380 | 2.853 |
| *Cntnap2*<sup>-/-</sup> female reversal errors | 5 | 15.20 errors | 1.789 | 0.8000 |
| **DeepLabCut Keypoint-MoSeq Open Field** |  |  |  |  |
| *Cntnap2*<sup>+/+</sup> male distance traveled | 7 | 272 m | 6112 | 2310 |
| *Cntnap2*<sup>-/-</sup> male distance traveled | 5 | 232 m | 4876 | 2181 |
| *Cntnap2*<sup>+/+</sup> female distance traveled | 6 | 202 m | 1604 | 654.9 |
| *Cntnap2*<sup>-/-</sup> female distance traveled | 6 | 227 m | 4651 | 1899 |
| *Cntnap2*<sup>+/+</sup> male number of grooming bouts (60 min.) | 7 | 324.7 bouts | 166.1 | 62.77 |
| *Cntnap2*<sup>-/-</sup> male number of grooming bouts (60 min.) | 5 | 411.6 bouts | 131.6 | 58.84 |
| *Cntnap2*<sup>+/+</sup> female number of grooming bouts (60 min.) | 6 | 361.0 bouts | 92.18 | 37.63 |
| *Cntnap2*<sup>-/-</sup> female number of grooming bouts (60 min.) | 6 | 534.0 bouts | 206.9 | 84.47 |

## Experimental procedures

### Mice

All animal procedures were conducted in accordance with protocols approved by the University of California, Berkeley Institutional Animal Care and Use Committee (IACUC) and Office of Laboratory Animal Care (OLAC) (AUP-2016-04-8684-3). *Cntnap2^−/−^* mice and littermate *Cntnap2^+/+^* controls with the following alleles were utilized for each experiment.

**Table 1.** Summary of mouse lines used.

| Experiment | Mouse line | Allele 1<br>(reference;<br>JAX strain #) | Allele 2<br>(reference;<br>JAX strain #) | Allele 3<br>(reference;<br>JAX strain #) | Allele 4<br>(reference;<br>JAX strain #) |
| --- | --- | --- | --- | --- | --- |
| Corticostriatal transmission<br>(Fig. 1) | *Cntnap2*;D1-tdTomato;Thy1-ChR2 | *Cntnap2*<br>(Poliak et al., 2003);<br>#017482) | *Drd1a*-tdTomato<br>(Ade et al., 2011);<br>#016204) | *Thy1*-ChR2-YFP<br>(Arenkiel et al., 2007);<br>#007612) |  |
| Broad inhibition<br>(Fig. 2A-E) | *Cntnap2*;D1-tdTomato | *Cntnap2*<br>(Poliak et al., 2003);<br>#017482) | *Drd1a*-tdTomato<br>(Ade et al., 2011);<br>#016204) |  |  |
| PV-specific inhibition<br>(Fig. 2F-J) | *Cntnap2*;D1-tdTomato;PV-Cre;Ai32 | *Cntnap2*<br>(Poliak et al., 2003);<br>#017482) | *Drd1a*-tdTomato<br>(Ade et al., 2011);<br>#016204) | *Pvalb*-Cre<br>(Hippenmeyer et al., 2005);<br>#017320) | Ai32<br>(Madisen et al., 2012);<br>#012569) |
| SPN intrinsic excitability<br>(Fig. 3, S4) | *Cntnap2*;D1-tdTomato | *Cntnap2*<br>(Poliak et al., 2003);<br>#017482) | *Drd1a*-tdTomato<br>(Ade et al., 2011);<br>#016204) |  |  |
| SPN intrinsic excitability in the presence of α-DTX<br>(Fig. 4, S5, S6) | *Cntnap2*;D1-tdTomato | *Cntnap2*<br>(Poliak et al., 2003);<br>#017482) | *Drd1a*-tdTomato<br>(Ade et al., 2011);<br>#016204) | | |
| Behavior experiments*<br>(Fig. 5-7, S7) | *Cntnap2*;D1-tdTomato | *Cntnap2*<br>(Poliak et al., 2003);<br>#017482) | *Drd1a*-tdTomato<br>(Ade et al., 2011);<br>#016204) |  |  |
| Spine analysis, PV cell counting | *Cntnap2*;D1-tdTomato | *Cntnap2* | *Drd1a*-tdTomato |  |  |
| (Fig. S1, S3A-H) |  | (Poliak et al., 2003); #017482) | (Ade et al., 2011); #016204) |  |  |
| PV intrinsic excitability (Fig. S2) | *Cntnap2*;PV-Cre;Ai9 | *Cntnap2* (Poliak et al., 2003); #017482) | *Pvalb*-Cre (Hippenmeyer et al., 2005); #017320) | Ai9 (Madisen et al., 2010); #007909) |  |
| PV in situ, western blot (Fig. S3I-N) | *Cntnap2* | *Cntnap2* (Poliak et al., 2003); #017482) |  |  |  |

\*Littermate animals both positive and negative for D1-tdTomato were used in behavior experiments

Mice were group housed on a 12 h light/dark cycle (dark cycle 9:00 AM – 9:00 PM) and given ad libitum access to standard rodent chow and water. Both male and female animals were used for experimentation. The ages, sexes, and numbers of mice used for each experiment are indicated in the respective method details and figure legends. All mice used for experiments were heterozygous or hemizygous for the *Drd1a*-tdTomato, *Thy1*-ChR2-YFP, PV-Cre, Ai32, or Ai9 transgenes to avoid potential physiological or behavioral alterations.

### Electrophysiology

Mice (P50-60) were briefly anesthetized with isoflurane and perfused transcardially with ice-cold ACSF (pH = 7.4) containing (in mM): 127 NaCl, 25 NaHCO3, 1.25 NaH2PO4, 2.5 KCl, 1 MgCl2, 2 CaCl2, and 25 glucose, bubbled continuously with carbogen (95% O_2_ and 5% CO_2_). Brains were rapidly removed and coronal slices (275 μm) were cut on a VT1000S vibratome (Leica) in oxygenated ice-cold choline-based external solution (pH = 7.8) containing (in mM): 110 choline chloride, 25 NaHCO3, 1.25 NaHPO4, 2.5 KCl, 7 MgCl2, 0.5 CaCl2, 25 glucose, 11.6 sodium ascorbate, and 3.1 sodium pyruvate. Slices were recovered in ACSF at 36°C for 15 min and then kept at room temperature (RT) before recording. Recordings were made with a MultiClamp 700B amplifier (Molecular Devices) at RT using 3-5 MOhm glass patch electrodes (Sutter, #BF150-86-7.5). Data were acquired using ScanImage software, written and maintained by Dr. Bernardo Sabatini (https://github. com/bernardosabatini/ SabalabAcq). Traces were analyzed in Igor Pro (Wavemetrics). Recordings with a series resistance > 25 MOhms or holding current above 200 pA were rejected. Passive properties were calculated using the double exponential curve fit of the average of five –5 mV, 100 ms long pulse steps applied at the beginning of every experiment.

#### Current-clamp recordings

Current clamp recordings were made using a potassium-based internal solution (pH = 7.4) containing (in mM): 135 KMeSO4, 5 KCl, 5 HEPES, 4 Mg-ATP, 0.3 Na-GTP, 10 phosphocreatine, and 1 EGTA. For corticostriatal excitability experiments, optogenetic stimulation consisted of a full-field pulse of blue light (470 nm, 0.5 ms pulse width, CoolLED) through a 63x objective (Olympus, LUMPLFLN60XW). Light power was linear over the range of intensities tested. No synaptic blockers were included. For intrinsic excitability experiments (SPN, PV interneuron, and SPN + α-DTX experiments), NBQX (10 μM, Tocris, #1044), CPP (10 μM, Tocris, #0247) and picrotoxin (50 μM, Abcam, #120315) were added to the external solution to block synaptic transmission. For Kv1.2 inhibition experiments, α-DTX (100nM, Alomone Labs, #D-350) was added to the external solution. Control recordings in the absence of α-DTX were performed on slices prior to drug application or on fresh slices after drug washout in alternating order across recording days. Bovine serum albumin (BSA, 0.005%, Sigma, #A7030) was included in both control and α-DTX-containing external solutions to minimize nonspecific binding. One second depolarizing current steps were applied to induce action potentials. No holding current was applied to the membrane.

#### Voltage-clamp recordings

Voltage-clamp recordings were made using a cesium-based internal solution (pH = 7.4) containing (in mM): 120 CsMeSO4, 15 CsCl, 10 TEA-Cl, 8 NaCl, 10 HEPES, 1 EGTA, 5 QX-314, 4 Mg-ATP, and 0.3 Na-GTP. Recordings were acquired with the amplifier Bessel filter set at 3 kHz. Corticostriatal synaptic stimulation experiments to measure evoked EPSCs were performed in picrotoxin (50 μM), and optogenetic stimulation consisted of a full-field pulse of blue light (470 nm, 0.15 ms pulse width) through a 63x objective. To record AMPAR-mediated EPSCs, cells were held at –70mV; to record NMDAR-mediated EPSCs, cells were held at +40mV. Synaptic stimulation experiments to measure evoked IPSCs were performed in NBQX (10 μM) and CPP (10 μM). For electrically evoked IPSCs, a concentric bipolar stimulating electrode (FHC, #30214) was placed in dorsal striatum, roughly 200 μm medial to the recording site in dorsolateral striatum, and a 0.15 ms stimulus was applied. For PV-interneuron optically evoked IPSCs, a full-field pulse of blue light (470 nm, 0.15 ms pulse width) was applied through a 63x objective at the recording site. All evoked IPSC experiments were recorded with cells held at –70mV.

### Dendritic imaging and spine analysis

Neonatal (P1-3) *Cntnap2^−/−^*;D1-tdT and *Cntnap2^+/+^*;D1-tdT mice were cryoanesthetized and injected bilaterally with 200 nL AAV1.hSyn.eGFP.WPRE.bGH (Penn Vector Core, #p1696 (Keaveney et al., 2018)), diluted 1:75 in saline to achieve sparse transduction. Injections were targeted to the dorsal striatum, with coordinates approximately 1.3 mm lateral to midline, 2.0 mm posterior to bregma, and 1.5 mm ventral to the head surface. At P50-60, mice were anesthetized with isoflurane and transcardial perfusion was performed with 10 mL of 1x PBS followed by 10 mL of ice cold 4% PFA (EMS, #15710-S) in 1x PBS. Brains were post-fixed in 4% PFA in 1x PBS overnight at 4° C. 80 μm coronal sections were made using a freezing microtome (American Optical, AO 860) and stored in 1x PBS at 4° C. Sections were blocked for 1 hour at RT in BlockAid (ThermoFisher, #B10710) and incubated for 48 hours with gentle shaking at 4° C with antibodies against GFP (1:2500, Abcam, #13970) and RFP (1:1000, Rockland (VWR, #600-401-379) diluted in PBS-Tx (1x PBS with 0.25% Triton X-100 (Sigma, #T8787). Sections were washed 3 x 10 min in PBS-Tx and incubated with gentle shaking for 1 hour at RT with Alexa Fluor 488 and 546 secondary antibodies (1:500, Invitrogen, #A11039, #A11035). Sections were washed 3 x 10 min in 1x PBS and mounted onto SuperFrost slides (VWR, #48311-703) using VECTASHIELD HardSet Antifade Mounting Medium (Vector Laboratories, #H-1400-10). Z stack images of individual dendrites were taken on a confocal microscope (Olympus FLUOVIEW FV3000) with a 60x oil immersion objective (Olympus #1-U2B832) at 2.5x zoom with a step size of 0.4 μm and deconvoluted using Olympus CellSens software. To quantify spine density, dendrites and spines were reconstructed using the FilamentTracer module in Imaris software (Oxford Instruments). The spine density of each dendrite was calculated using Imaris. Dendrites analyzed varied in total length, but excluded the most proximal and distal portions of the dendrite.

### Brain sectioning and immunohistochemistry

Adult mice were perfused as above and brains were post-fixed with 4% paraformaldehyde overnight, then sectioned coronally at 30 μm. For immunohistochemistry, individual wells of sections were washed for 3 x 5 min with 1x PBS, then blocked for 1 hour at RT with BlockAid blocking solution. Primary antibodies diluted in PBS-Tx were added and tissue was incubated for 48 hours with gentle shaking at 4° C. Sections were then washed 3 x 10 min with PBS-Tx. Secondary antibodies diluted 1:500 in PBS-Tx were added and incubated with gentle shaking for 1 hour at RT. Sections were washed 3 x 10 min in 1x PBS. Sections were mounted onto SuperFrost slides (VWR, #48311-703) and coverslipped with VECTASHIELD HardSet with DAPI (Vector Laboratories, #H-1500-10) or VECTASHIELD HardSet Antifade Mounting Medium (Vector Laboratories, #H-1400-10). The following antibodies were used: mouse anti-PV (1:1000, Sigma, #P3088), rabbit anti-PV (1:1000, Abcam, #11427), anti-RFP (1:500, Rockland, #600-401-379), Alexa Fluor 405, 488 and 546 conjugated secondary antibodies (1:500, Invitrogen, #A-31553, #A-11001, #A-11003, and #A-11035).

#### PV cell counting

To count PV+ interneurons, Z-stack images of immunostained striatal sections were taken on a confocal microscope (Olympus FLUOVIEW FV3000) with a 10x or 20x objective (Olympus # 1-U2B824 or Olympus # 1-U2B825) and step size of 2 μm. For quantification, image stacks were Z-projected to maximum intensity using Fiji (ImageJ) and cropped to a 400 μm x 400 μm image in anatomically matched sections of the DLS. All PV-expressing cells within this region were counted using the ROI manager tool in ImageJ. Designation of a cell as PV positive was determined by the experimenter and consistently maintained across animals. Experimenter was blind to genotype and ROIs were made on the DAPI channel to avoid selecting regions based on PV expression. To quantify bulk PV fluorescence, ROIs were manually defined in ImageJ using the Freehand tool to cover as much of the DLS as possible, and mean fluorescence intensity was measured. To quantify individual cell PV fluorescence, ROIs were manually defined around every PV positive cell in the previously drawn DLS ROI using the Freehand tool, and mean fluorescence intensity was measured.

### Western Blot

Adult mice (P48-55) were deeply anesthetized with isoflurane and decapitated. Brains were rapidly dissected and 1.5 mm dorsal striatum punches (Biopunch, Ted Pella, #15111-15) were collected from both hemispheres, flash-frozen in liquid nitrogen, and stored at −80° C. On the day of analysis, frozen samples were sonicated (QSonica Q55) until homogenized in 200 μl lysis buffer containing 1% SDS in 1x PBS with Halt phosphatase inhibitor cocktail (Thermo Fisher Scientific, #PI78420) and Complete mini EDTA-free protease inhibitor cocktail (Roche, #4693159001). Sample homogenates were then boiled on a heat block at 95° C for 5 min and allowed to cool to RT. Total protein content was determined using a BCA assay (Thermo Fisher Scientific, #23227). Following the BCA assay, protein homogenates were mixed with 4x Laemmli sample buffer (BioRad, #161-0747). 12.5μg of total protein per sample were then loaded onto 12% Criterion TGX gels (BioRad, #5671044) and run at 65 V. Proteins were transferred to a PVDF membrane (BioRad, #1620177) at 11 V for 14 hours at 4° C using the BioRad Criterion Blotter (BioRad, #1704070). Membranes (BioRad, #1620177) were briefly reactivated in methanol and rinsed in water 3x. After rinsing, membranes were blocked in 5% milk in 1x TBS with 1% Tween (TBS-Tween) for 1 hour at RT before being incubated with primary antibodies diluted in 5% milk in TBS-Tween overnight at 4° C. The following day, after 3 x 10 min washes with TBS-Tween, membranes were incubated with secondary antibodies for 1 hour at RT. Following 6 × 10 min washes, membranes were incubated with chemiluminescence substrate (PerkinElmer #NEL105001EA) for 1 min and exposed to Amersham Hyperfilm ECL (VWR, #95017-661).

Bands were quantified by densitometry using ImageJ software. GAPDH was used to normalize protein content and data are expressed as a percentage of control within a given experiment. The following antibodies were used: anti-Caspr2 (1:5000, Abcam, #153856), anti-PV (1:2500, Abcam, #11427), anti-GAPDH (1:5000, Cell Signaling, #51745S), and anti-rabbit goat HRP conjugate (1:5000, BioRad, #1705046).

### In situ hybridization

Fluorescent in situ hybridization was performed to quantify *Pvalb* mRNA expression in the striatum of *Cntnap2^+/+^* and *Cntnap2^−/−^* mice. Mice were briefly anesthetized with isoflurane and brains were harvested, flash-frozen in OCT mounting medium (Thermo Fisher Scientific, #23-730-571) on dry ice and stored at –80° C for up to 6 months. 16 µm sections were collected using a cryostat (Thermo Fisher Scientific, Microm HM 550), mounted directly onto Superfrost Plus glass slides (VWR, #48311-703) and stored at –80° C for up to 6 months. In situ hybridization was performed according to the protocols provided with the RNAscope Multiplex Fluorescent Reagent Kit (ACD, #323100). *Drd1a* mRNA was visualized with a probe in channel 2 (ACD, #406491-C2) and *Pvalb* mRNA in channel 3 (ACD, #421931-C3). After incubation, sections were secured on slides using ProLong Gold Antifade Mountant with DAPI (Invitrogen, P36935) and 60 x 24 mm rectangular glass coverslips (VWR, #16004-096). Sections were imaged on an Olympus FluoView 3000 confocal microscope using a 10x objective with 1.5x zoom and a step size of 2 µm. *Pvalb*-expressing cells were quantified across the entire striatum using the ROI manager tool in ImageJ. A cell was considered *Pvalb* positive if over 50% of the cell contained fluorescent puncta when compared to the DAPI channel. Experimenter was blind to genotype.

### Behavioral analysis

All behavior studies were carried out in the dark phase of the light cycle under red lights (open field) or white lights (marble burying, holeboard, rotarod, and four choice reversal learning). Mice were habituated to the behavior testing room for at least 30 min prior to testing. Mice were given at least one day between different tests. All behavior equipment was cleaned between each trial and mouse with 70% ethanol and rinsed in diluted soap followed by water at the end of the day. If male and female mice were to be tested on the same day, male mice were run first then returned to the housing room, after which all equipment was thoroughly cleaned prior to bringing in female mice for habituation. Behavioral tests were performed with young adult male and female mice (7-11 weeks old). The experimenter was blind to genotype throughout the testing and scoring procedures.

#### Open field assay

Exploratory behavior in a novel environment and general locomotor activity were assessed by a 60 min session in an open field chamber (40 cm L x 40 cm W x 34 cm H) made of transparent plexiglass. Horizontal infrared photobeams (Stoelting, 60001-02A) were positioned to detect rearing. The mouse was placed in the bottom right-hand corner of the arena and behavior was recorded using an overhead camera and analyzed using ANY-maze software (Stoelting). An observer manually scored self-grooming behavior during the first 20 minutes of the test. A grooming bout was defined as an unbroken series of grooming movements, including licking of body, paws, or tail, as well as licking of forepaws followed by rubbing of face with paws.

#### Open field assay with DeepLabCut Keypoint-MoSeq analysis

Mice were placed in the open field arena and video recorded with a monochrome camera (FLIR Grasshopper 3, GS3-U3-41C6NIR-C) and a 16 mm wide angle lens (Kowa, LM16HC) placed above the arena from a height of 50 cm. To extract the body part (keypoint) coordinates from the video recordings, DeepLabCut (DLC) 2.3.4 (Mathis, et al. 2018; Nath, et al. 2019) was used. Fourteen body parts including nose, head, left ear, right ear, left forelimb, right forelimb, spine 1, spine 2, spine 3, left hindlimb, right hindlimb, tail 1, tail 2, and tail 3 were manually labeled on a small subset of the video frames. A DLC model was then trained using the annotated frames to label those 14 body parts for all videos recorded. The total distance traveled, and number of center entries were calculated using the coordinate of bodypart tail 1.

Discrete behavior syllables were extracted using Keypoint-MoSeq 0.4.4 (Weinreb, et al. 2023). Syllable usage and transition data were obtained using built-in functions of the Keypoint-MoSeq package. Decoding and entropy analysis were performed using customized Python 3.9 script. Code available upon request in Github. Entropy was calculated using the following equation, where *u*_i_ denotes the frequency of the syllable *i* and *p*_i,j_denotes the transition probability from syllable *i* to syllable *j*.:

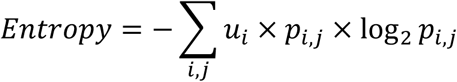

#### Marble burying assay

The marble burying assay was used to test for repetitive behavior. 20 black marbles were arranged in an orderly 4 x 5 grid on top of 5 cm of clean corn cob bedding in a standard mouse cage. Overhead room lights were on and white noise was played to induce mild stress. Mice were placed in the cage with the marbles for 30 minutes. The number of unburied marbles (>50% exposed) was recorded after the session.

#### Holeboard assay

The holeboard assay was used to measure exploratory and repetitive behavior. The holeboard apparatus consisted of a smooth, flat, opaque gray plastic platform, suspended 10 cm from the base by four plastic pegs in each corner. The board contained 16 evenly spaced 2 cm diameter holes and was surrounded by a 30 cm high clear plastic square encasing. During testing, mice were placed into the center of the holeboard. Mice explored the board for 10 minutes while video was recorded from both an above and side-view camera. Videos were used post-hoc to manually count and map the number of nose pokes made during the task. Nose pokes were defined as the mouse’s nose passing through the board barrier when viewed through the side-view camera.

#### Accelerating rotarod assay

The accelerating rotarod test was used to examine motor coordination and learning.

Mice were trained on a rotarod apparatus (Ugo Basile, #47650) for four consecutive days. Three trials were completed per day with a 5 min break between trials. The rotarod was accelerated from 5-40 revolutions per minute (rpm) over 300 s for trials 1-6 (days 1 and 2), and from 10-80 rpm over 300 s for trials 7-12 (days 3 and 4). On the first testing day, mice were first acclimated to the apparatus by being placed on the rotarod rotating at a constant 5 rpm for 60 s and returned to their home cage for 5 min prior to starting trial 1. Latency to fall, or to rotate off the top of the rotarod barrel, was measured by the rotarod stop-trigger timer.

#### Four choice odor-based reversal learning test

The four-choice odor-based reversal learning test was used to assess learning and cognitive flexibility. Animals were food restricted for 6 days in total, with unrestricted access to drinking water, and maintained at 90-95% of ad lib feeding body weight. Food was given at the end of the day once testing was completed. Food restriction and introduction to Froot Loop cereal piece (Kellogg’s, Battle Creek, MI) began 48 hours before pre-training. The four-choice test was performed in a custom-made square box (30.5 cm L × 30.5 cm W × 23 cm H) constructed of clear acrylic. Four internal walls 7.6 cm wide partially divided the arena into four quadrants. A 15.2 cm diameter removable cylinder fit in the center of the maze and was lowered between trials (after a digging response) to isolate the mouse from the rest of the maze. Odor stimuli were presented mixed with wood shavings in white ceramic pots measuring 7.3 cm in diameter and 4.5 cm deep. All pots were sham baited with a piece of Froot Loop cereal secured underneath a mesh screen at the bottom of the pot. This was to prevent mice from using the odor of the Froot Loop to guide their choice. The apparatus was cleaned with 2.5% acetic acid followed by water and the pots were cleaned with 70% ethanol followed by water between mice. The apparatus was cleaned with diluted soap and water at the end of each testing day.

On the first habituation day of pre-training (day 1), animals were allowed to freely explore the testing arena for 30 min and consume small pieces of Froot Loops placed inside empty pots positioned in each of the four corners. On the second shaping day of pre-training (day 2), mice learned to dig to find cereal pieces buried in unscented coarse pine wood shavings (Harts Mountain Corporation, Secaucus, NJ). A single pot was used and increasing amounts of unscented wood shavings were used to cover each subsequent cereal reward. The quadrant containing the pot was alternated on each trial and all quadrants were rewarded equally. Trials were untimed and consisted of (in order): two trials with no shavings, two trials with a dusting of shavings, two trials with the pot a quarter full, two trials with the pot half full, and four trials with the cereal piece completely buried by shavings. The mouse was manually returned to the center cylinder between trials.

On the days for odor discrimination (day 3, acquisition) and reversal (day 4), wood shavings were freshly scented on the day of testing. Anise extract (McCormick, Hunt Valley, MD) was used undiluted at 0.02 ml/g of shavings. Clove, litsea, and eucalyptus oils (San Francisco Massage Supply Co., San Francisco, CA) were diluted 1:10 in mineral oil and mixed at 0.02 ml/g of shavings. Thymol (thyme; Alfa Aesar, A14563) was diluted 1:20 in 50% ethanol and mixed at 0.01 ml/g of shavings. During the discrimination phase (day 3), mice had to discriminate between four pots with four different odors and learn which one contained a buried food reward. Each trial began with the mouse confined to the start cylinder. Once the cylinder was lifted, timing began, and the mouse could freely explore the arena until it chose to dig in a pot. Digging was defined as purposefully moving the shavings with both front paws. A trial was terminated if no choice was made within 3 min and recorded as omission. Criterion was met when the animal completed eight out of ten consecutive trials correctly. The spatial location of the odors was shuffled on each trial. The rewarded odor during acquisition was anise.

The first four odor choices made during acquisition were analyzed to determine innate odor preference by the percentage of choices for a given odor: *Cntnap2^+/+^* mice: 60% thyme, 25% anise, 12.5% clove, and 2.5% litsea. *Cntnap2^−/−^* mice: 47.5% thyme, 45% anise, 7.5% clove, 0% litsea. We note that both *Cntnap2^+/+^* and *Cntnap2^−/−^* mice exhibited the strongest innate preference for thyme, an unrewarded odor. There were no significant differences in innate odor preference.

The reversal phase of the task was carried out on day 4. Mice first performed the task with the same rewarded odor as the discrimination day to ensure they learned and remembered the task. After reaching criterion on recall (eight out of ten consecutive trials correct), the rewarded odor was switched, and mice underwent a reversal learning test in which a previously unrewarded odor (clove) was rewarded. A novel odor (eucalyptus) was also introduced, which replaced thyme. Perseverative errors were choices to dig in the previously rewarded odor that was no longer rewarded. Regressive errors were choosing the previously rewarded odor after the first correct choice of the newly rewarded odor. Novel errors were choices to dig in the pot with the newly introduced odor (eucalyptus). Irrelevant errors were choices to dig in the pot that had never been rewarded (litsea). Omissions were trials in which the mouse failed to make a digging choice within 3 min from the start of the trial. Total errors were the sum of perseverative, regressive, irrelevant, novel, and omission errors. Criterion was met when the mouse completed eight out of ten consecutive trials correctly. The spatial location of the odors was shuffled on each trial.

### Quantification and statistical analysis

Experiments were designed to compare the main effect of genotype. The sample sizes were based on prior studies and are indicated in the figure legend for each experiment. Whenever possible, quantification and analyses were performed blind to genotype. GraphPad Prism version 10 was used to perform statistical analyses. The statistical tests and outcomes for each experiment are indicated in the respective figure legend. Two-tailed unpaired t tests were used for comparisons between two groups. For data that did not pass the D’Agostino & Pearson normality test, a Mann-Whitney test was used. Two-way ANOVAs or mixed effects models were used to compare differences between groups for experiments with two independent variables. Statistical significance was defined in the figure panels as follows: *p < 0.05, **p < 0.01, ***p < 0.001.

## Acknowledgements

This work was supported by Simons Foundation Autism Research Initiative (SFARI) research grant #514428 to H.S.B. and NIH fellowship #F31NS124499 to K.R.C.

